# Dynamic genome-based metabolic modeling of the predominant cellulolytic rumen bacterium *Fibrobacter succinogenes* S85

**DOI:** 10.1101/2022.10.18.512662

**Authors:** Ibrahim Fakih, Jeanne Got, Carlos Eduardo Robles-Rodriguez, Anne Siegel, Evelyne Forano, Rafael Muñoz-Tamayo

## Abstract

*Fibrobacter succinogenes* is a cellulolytic predominant bacterium that plays an essential role in the degradation of plant fibers in the rumen ecosystem. It converts cellulose polymers into intracellular glycogen and the fermentation metabolites succinate, acetate, and formate. We developed dynamic models of *F. succinogenes* S85 metabolism on glucose, cellobiose, and cellulose on the basis of a network reconstruction done with the Automatic Reconstruction of metabolic models (AuReMe) workspace. The reconstruction was based on genome annotation, 5 templates-based orthology methods, gap-filling and manual curation. The metabolic network of *F. succinogenes* S85 comprises 1565 reactions with 77% linked to 1317 genes, 1586 unique metabolites and 931 pathways. The network was reduced using the NetRed algorithm and analyzed for computation of Elementary Flux Modes (EFMs). A yield analysis was further performed to select a minimal set of macroscopic reactions for each substrate. The accuracy of the models was acceptable in simulating *F. succinogenes* carbohydrate metabolism with an average coefficient of variation of the Root mean squared error of 19%. Resulting models are useful resources for investigating the metabolic capabilities of *F. succinogenes* S85, including the dynamics of metabolite production. Such an approach is a key step towards the integration of omics microbial information into predictive models of the rumen metabolism.

## 1. Introduction

The rumen microbiota plays an essential role in ruminant nutrition by breaking down and fermenting plant-based feed, transforming it into a source of energy and protein for the host. The rumen microbiota is composed of a very diverse community of prokaryotes (bacteria and archaea) and eukaryotes (protozoa and fungi) which concur to the degradation and fermentation of the feed components, and particularly complex fibrous substrates that cannot be digested by the host. Rumen bacteria, fungi and protozoa participate to the degradation of the plant cell wall lignocellulose (1), producing a large array of enzymes and various enzymatic systems to deconstruct the intricate chemical structure of plant biomass (2). Among them, cellulose degraders have been particularly studied for decades, because cellulose is the most degradation-resistant polysaccharide in plants, and it represents an abundant renewable resource on earth (3). Within cellulolytic bacteria, *Fibrobacter succinogenes* has been particularly studied (2). *F. succinogenes* is found in large numbers in ruminants fed high fiber diets (4), and is present in the rumen of farm but also wild ruminant species from many geographical regions worldwide (5). It has been quantified in higher levels in bovines compared to deer, sheep, or camelids, suggesting that it may play an essential role in plant fiber degradation in cattle. *F. succinogenes* belongs to the *Fibrobacteres* phylum which also comprises the species *F. intestinalis*, mainly isolated from the feces of ruminant and non-ruminant animals (6).

The strain *F. succinogenes* S85 has been isolated from a bovine rumen a long time ago (7, 8), and is the most studied strain of the species. For efficient plant cell wall degradation, *F. succinogenes* adheres closely to the substrate and produces specific cellulose-binding proteins and possibly also pili to mediate its adhesion (9–11). *F. succinogenes* is considered as particularly efficient in the hydrolysis of crystalline cellulose, and it degrades at the same rate amorphous and crystalline regions of wheat straw cellulose (12). Cellulose is degraded into cellodextrins, cellobiose and glucose, and *F. succinogenes* was shown to be a very effective competitor for cellodextrin utilization (13). The bacterium is also able to synthesize and efflux oligosaccharides that may be used by other rumen bacteria through cross-feeding (12, 14). Given all these properties, it may be interesting to promote *F. succinogenes* populations in the rumen of cattle to improve degradation of recalcitrant substrates and their utilization by the rumen microbiota.

The analysis of the *F. succinogenes* S85 genome showed that it consists of approximately 3.84 Mbp with a GC content of 48%, and that it contains a high number of genes (134) encoding carbohydrate-active enzymes (CAZymes) (10). The genome analysis also confirmed that despite its ability to degrade xylans (15), *F. succinogenes* cannot use xylose because it lacks the sugar transporter and phosphorylation system (16). This species is thus a cellulose specialist using cellulose and its degradation products as its sole energy source. Glucose and cellobiose are fermented mainly through the EMP pathways into succinate as major final product, followed by acetate, formate and CO_2_. *F. succinogenes* is able to store intracellular glycogen which can represent up to 70% of the dry weight of the bacterium (17). This storage could allow bacteria to remain in the rumen in the absence of metabolizable substrates (18), but the intracellular glycogen is simultaneously stored and degraded, suggesting a futile cycling (19). *F. succinogenes* uses ammonia as the sole source of nitrogen, and several steps in the ammonia assimilation pathway have been identified (20). In addition to its interest for ruminant nutrition, *F. succinogenes* has also received much attention from the biotechnology sector (21). Firstly, because this species produces an original cellulolytic system, whose organization is still not well understood, and includes membrane vesicles as vehicles of CAZymes (22). Deciphering this system could help in the design of novel Consolidated Bioprocessing (CBP) for the production of cost-effective and sustainable lignocellulosic biofuels (9). Secondly, the capacity of *F. succinogenes* to transform lignocellulosic material into succinate may also be of interest because succinic acid could be used as a platform molecule (23, 24). However, increasing bacterial product yield or maximizing the production of powerful lignocellulose degradation enzymes is dependent on detailed knowledge of metabolic pathways for microbial engineering processes (25). An efficient way of deciphering a bacterium metabolic network and identifying possible bottlenecks in the production of metabolites is *via* genome-scale metabolic models (GEM). A GEM is a mathematical representation of a metabolic network that allows the study of genotype-phenotype relationships (26) and facilitates the prediction of multiscale phenotypes (27). For a genome-sequenced microorganism, a GEM is defined by a stoichiometry matrix that links metabolites to the collection of reactions that occur in the organisms according to evidences about genes catalyzing the reactions. The resulting metabolic network can be further analyzed using methods such as flux balance analysis (FBA) (28–30). However, the FBA approach does not allow to predict the dynamics of metabolites concentrations. In parallel, kinetic modeling approaches allow to represent the dynamics of metabolites of interest by deriving mass balance equations (31). Kinetic models are built following a macroscopic representation of the metabolism with a reduced set of macroscopic reactions which are often selected from documented literature. Kinetic models rarely integrated microbial genomic information. The objective of this work was to develop dynamic metabolic models (DMM) to represent the metabolism of glucose, cellobiose and cellulose by of *F. succinogenes*. These DMM integrate microbial genomic knowledge from the reconstruction of a GEM of *F. succinogenes*.

## 2. Materials and Methods

### 2.1. Culture conditions and sample preparation

*F. succinogenes* strain S85 (ATCC 19169) was grown in triplicates in a chemically defined medium (19) with 3 g.L^-1^ glucose, cellobiose, or filter paper cellulose. The cultures were grown at 39°C using Hungate tubes and the Hungate anaerobic cultivation technique (32). The bacterial growth on cellobiose or glucose was monitored by measuring the absorbance at 600 nm. The quantification of succinate was used to monitor growth on cellulose cultures (12, 33). After growth, cells were collected by centrifugation (10 000 rpm for 5 min at 4°C), and the supernatants were stored at −20°C for further analysis. The bacteria were washed with 2 mL of phosphate buffer (50 mM KH_2_PO_4_/K_2_HPO_4_ pH 7.0) and stored at −20°C.

### 2.2. Quantification of substrate consumption and metabolite production

Concentrations of succinate, acetate, formate, ammonia, and glucose were measured at six time points in culture supernatants, using Megazyme kits (K-SUCC 06/18, K-ACET 04/18, K-FORM 10/17, K-AMIAR 04/18, and K-GLUHK-220A, respectively) according to the manufacturer’s recommendations. Cellobiose consumption was estimated by quantification of the remaining reducing sugars in the culture medium using Miller’s method (34).

### 2.3. Metabolic network reconstruction

The metabolic reconstruction of *F. succinogenes* S85 was performed using the freely available workspace AuReMe (Automatic Reconstruction of Metabolic models) (35). AuReMe embeds existing tools as well as ad-hoc packages to reconstruct and handle GEMs. It uses (i) the outputs of the Pathway Tools software (36, 37) to perform annotation-based reconstruction, (ii) the OrthoFinder method (38, 39) to perform orthology-based reconstructions and (iii) the Meneco tool (40) to perform a gap-filling procedure. AuReMe relies on the PADMet library (Python library for hAndling metaData of METabolism, (35) to ensure the reproducibility of the workflow used to create a GEM, guaranty the interoperability between the different methods used, curate GEMs and export the GEM under several formats (SBML, matrix, wiki, rdf-like format). AuReMe also uses CobraPy (41), a Python package to analyze Flux Balance Analysis, and Flux Variability analysis (analysis of essential and blocked reactions). In Fig. 1, we summarize the steps of reconstruction of *F. succinogenes* S85 metabolic network.

**Figure 1.**
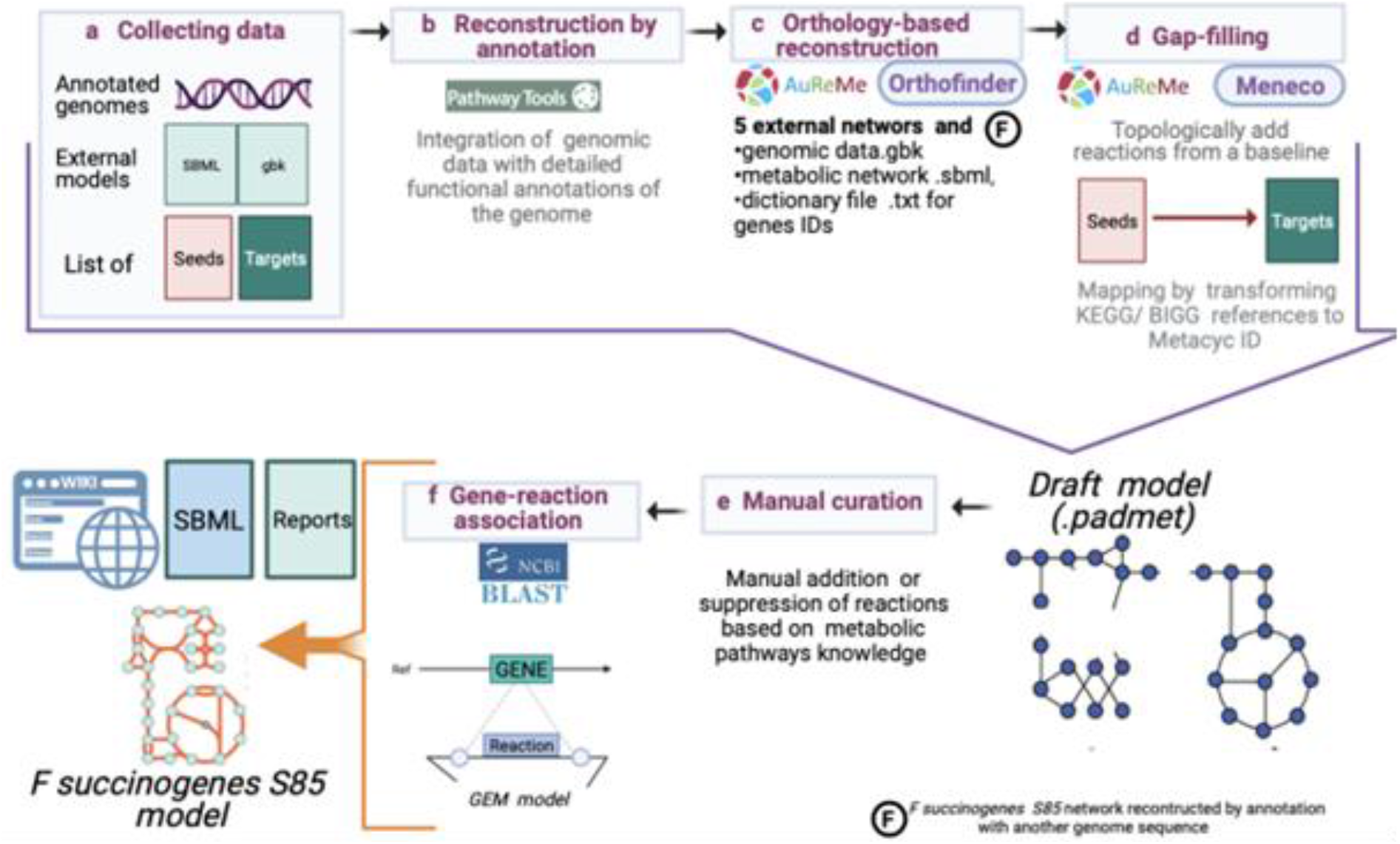
F. succinogenes S85 GEM reconstruction pipeline.

#### 2.3.1. Step 1 (collecting genomes and reference reaction data set)

The two complete annotated genomes of *F. succinogenes* S85 were downloaded respectively from https://www.ncbi.nlm.nih.gov/nuccore/CP001792.1 and https://www.ncbi.nlm.nih.gov/nuccore/NC_017448.1. Each *F. succinogenes* S85 genome contains 3.84 Mbp with respectively 3160 and 3174 genes identified.

The biomass reaction of *Escherichia coli* K-12 MG1655 (42) was adapted and used to build the metabolic network (Table S1 in Supplementary Material A).

The list of seeds (essential constituents of the culture medium to guarantee growth) was prepared based on the minimal medium composition needed for *F. succinogenes* growth (Table S2). The final products (targets) list was prepared according to our knowledge on the metabolism of this bacterium metabolism and network reconstruction needs (Table S2).

#### 2.3.2. Step 2 (generating draft models)

A first GEM was reconstructed according to the genome annotations via Pathway Tools using both complete genomes: NC_013410.1. and NC_017448. In parallel, orthology-based reconstruction GEMs we obtained using the GEMs of the following gut microbes: *Bacteroïdes thetaiotaomicron* (43), *Escherichia coli* K-12 MG1655 (42), *Faecalibacterium prausnitzii* A165 (29), *Bifidobacterium adolescentis* L2-32 (28) and *Lactobacillus plantarum* WCFS1 (44), and mapped to MetaCyc (45), thanks to the MetaNetX database (46). Finally, all GEMs obtained from the annotation and the orthology reconstruction steps were combined into a draft GEM with the PADMet library.

#### 2.3.3. Step 3 (gap filling)

To analyze and curate the GEM of *F. succinogenes* S85, we applied a gap-filling procedure. Here, a GEM is considered as a graph in which metabolites are nodes and reactions are the links between the nodes. In these analyses, stoichiometry is not considered. This procedure allows adding reactions to guarantee the production of specific metabolites according to a graph-expansion criterion. We used Meneco (MEtabolic NEtwork COmpletion, (40)) and the MeneTools package (MEtabolic NEtwork TOpological toOLS, (35, 47)) for this gap-filling step.

#### 2.3.4. Step 4 (manual curation and gene-reaction association)

The draft network was manually curated to find potential errors and filling gaps based on the phenotype and experimental data reported in the literature. Flux Balance Analysis (FBA) was used to reconstruct and validate models for maximizing the biomass reaction flux.

Some manually and gap-filled added reactions had no gene associated. All the gene sequences from other bacteria associated with these reactions were identified in NCBI using the reaction EC number. The corresponding protein sequences were aligned using BLAST (Basic Local Alignment Search Tool, (48)) to the *F. succinogenes* S85 translated genomes. The identified proteins with identity > 76% were associated with their corresponding reactions in the GEM. Reactions with no gene associated or with gene coding protein of lower similarity were retained in the model only when present in the *E. coli* K-12 MG1655 model (42).

### 2.4. Construction of a dynamic metabolic model

#### 2.4.1. Exploiting EFMs to derive a macroscopic dynamic metabolic model

The dynamics of metabolism can be described by the following generic differential equation **(1)** resulting from applying mass balance

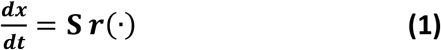

where ***x*** is the vector containing the concentrations of metabolites, which can be either intracellular (***x**_i_*) or extracellular (***x**_e_*). The vector ***r***(·) represents the reaction rates, which are function of the concentrations ***x*** and a parameter vector. The stoichiometric matrix ***S*** contains the stoichiometric matrices for intracellular (***S**_i_*) and extracellular ***S**_e_* metabolites. Under the assumption that intracellular metabolism operates at steady state, it follows that

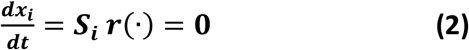

The vector of reaction rates that fulfil equation **(2)** are non-negative vectors contained in the null space of the stoichiometric matrix ***S***_i_. The space of admissible fluxes is a convex polyhedral cone. The generating vectors of the cone are called elementary flux modes (EFMs). Any steady-state flux distribution can be expressed as a non-negative linear combination of the EFMs. Biochemically, EFMs are independent minimal pathways of the metabolic network that can operate at steady state. Each EFM can be converted into a macroscopic reaction that connects extracellular substrates and products (49, 50). The identification of macroscopic reactions is the core of kinetic modeling. Once we find a set of macroscopic reactions to represent the metabolism of our microorganism of interest, we can derive the structure of a dynamic metabolic model.

For the *j*-th EFM ***e**_j_*, the macroscopic reaction *j* is obtained by the product ***S**_e_**e**_j_*. To calculate the EFMs of the network of *F. succinogenes*, we used the efmtool algorithm (51) of the MATLAB package CellNetAnalyzer (52) which is freely available at http://www2.mpi-magdeburg.mpg.de/projects/cna/cna.html Then, we proceeded to select a reduced number of EFMs using the yield analysis method by (53). The selected EFMs were expressed as macroscopic reactions to define further the kinetics in the dynamic metabolic modelling.

#### 2.4.2. Reduction of the GEM

The calculation of EFMs is restricted to medium-scale GEM (less than 350 reactions) (54). Hence, a complete EFM analysis of the network of *F. succinogenes* S85 is intractable. A reduction of the network is thus here proposed. Several methods for the reduction of GEM have been reported in the literature (55) which are mainly based on a fully functional core metabolic network that preserves a set of important moieties and capabilities from the full network. However, the selection of a subset of reactions might produce loses of information regarding parallel pathways that can be used to attain the same metabolic goal. In this work, we have selected another method called NetRed (56) which analyses flux vectors generated from the complete network (FBA) and computes a reduced network that holds the same flux distribution. NetRed is based on matrix algebra taking as inputs the stoichiometric matrix, a flux vector, the numerical flux values, and a list of protected metabolites, and it is implemented in the MATLAB COBRA toolbox (57). Some advantages of NetRed are the use of various flux vectors, the calculation of a single biomass reaction, and the simple conversion from the reduced network to the full network.

The reduction of the GEM followed several steps: (i) calculation of fluxes by FBA, (ii) carbon balancing, (iii) compacted lumped biomass reaction, (iv) re-calculation of fluxes by FBA, and (v) network reduction.

To keep the flexibility of the full network, the flux distribution of the GEM was calculated by FBA considering different objective functions and several input fluxes. The results were analyzed in terms of yields for which all the obtained fluxes were divided by the uptake flux of glucose, cellobiose or cellulose. Yield analysis allowed to verify the carbon balance of the network as well.

Further reduction of the network was achieved by computing a compact lumped biomass reaction, which was based on the pathways identified as essential for biomass production. This approach is like the construction of a core metabolism. In our case, however, we have only used this core for biomass allowing for other pathways to contribute as well to the production of metabolites needed for biomass. Finally, the obtained fluxes from the network with compacted biomass formulation and correct carbon balance have been introduced to NetRed to compute the final reduced network which was used to compute EFMs.

#### 2.4.3. Parameter identification

The model parameters were estimated from the *in vitro* experimental data (acetate, succinate, formate, ammonia and OD and substrate concentration when available) using the maximum likelihood approach implemented in the MATLAB toolbox IDEAS (58), which is freely available at http://genome.jouy.inra.fr/logiciels/IDEAS. The optimization uses the quasi-newton algorithm implemented in the MATLAB function *fminunc*. The dynamic model representing the fermentation of each substrate is defined by the kinetic rate function of substrate utilisation through the macroscopic reactions given by the EFMs. For glucose and cellobiose, we modelled the macroscopic reactions as Monod functions **(3)**

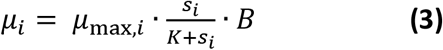

where *s_i_* and *B* are the molar concentrations of the substrate and biomass, *μ_i_* is the microbial growth rate for the reaction *i*, *μ*_max_, *i* is the maximal growth rate constant (h^-1^) and *K* is the substrate affinity Monod constant (mol/L). We set *K* to 9*10^-3^ M as in the rumen model developed by Muñoz-Tamayo et al (59) to avoid the known problems of high correlation between the Monod parameters when data are limited (60). Since cellulose is a particulate substrate, we modelled the macroscopic reactions using the Contois function **(4)** as proposed by Vavilin et al(61).

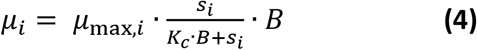

where *K_c_* is the half-saturation Contois constant. For each substrate, we selected initially the EFMs that correspond to the vertices of the polygon enclosing the yield spaces. A further reduction was implemented within the calibration procedure by adding a penalization coefficient in the cost function of the optimization to penalize large number of EFMs. To account for the death of microbial cells, we included a first-order kinetic rate with a death rate constant *k_d_* set to 8.33×10^-4^ as in (59). We also included a conversion factor to transform OD into biomass concentration (mol/L). We evaluated the model accuracy using the coefficient of variation of the Root mean squared error (CVRMSE).

## 3. Results

Following Open Science practices to promote accessibility and reproducibility (62), the metabolic network and mathematical models developed in this work are freely available at https://doi.org/10.5281/zenodo.7219865.

### 3.1 Description of the network

#### • Large-scale Genome reconstruction process

The two published genomes of the *Fibrobacter succinogenes S85* strain were used to identify potential reactions that could be present in the GEM of the bacterium. Genome annotation performed by Pathway Tools had detected 827 reactions (Fig. 2A) and 1112 metabolites. 817 reactions are common between the two available *F. succinogenes* S85 annotated genomes. Four and six reactions were specific to NC_013410 and NC_017448 genomes, respectively (Table S3 in Supplementary Material A), illustrating that the two genome sequences are not complete.

**Figure 2.**
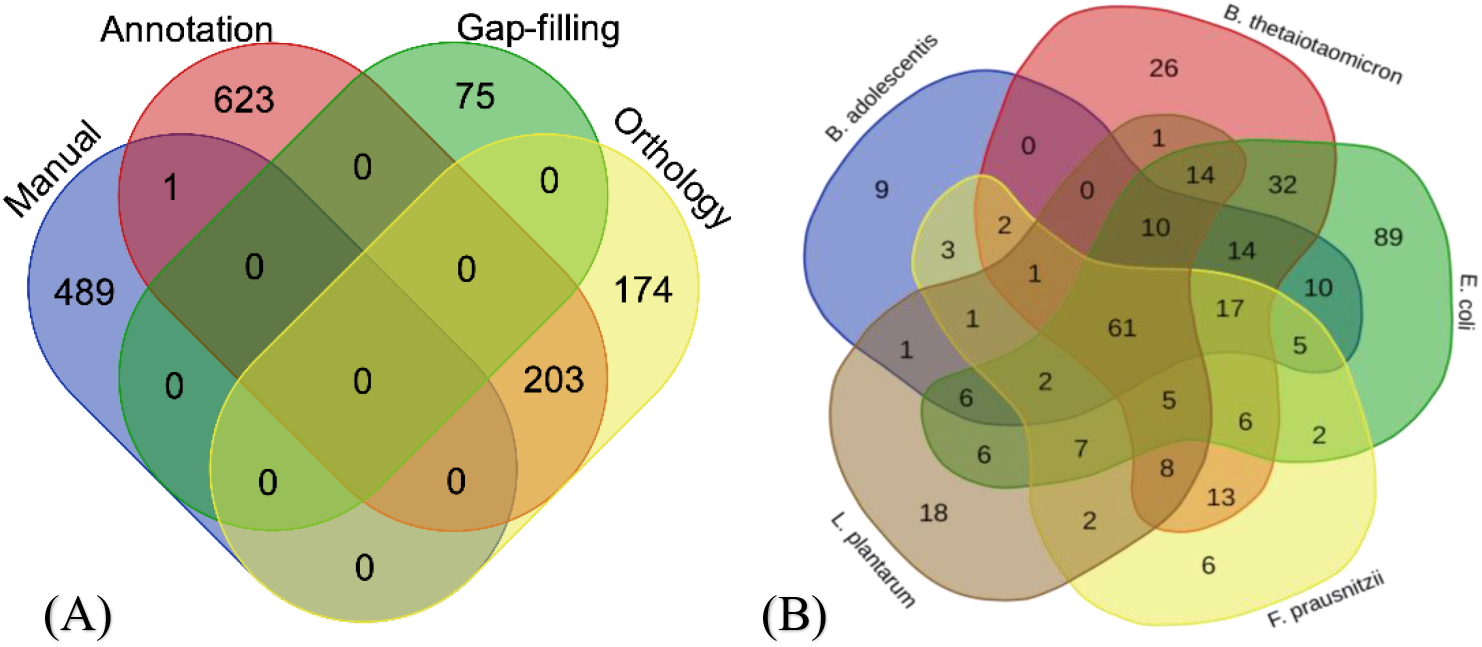
Venn diagram representing (A) distribution of reactions across different steps of final model reconstruction (B) distribution of reactions across various external metabolic models used for orthology-based model reconstruction.

#### • Orthology and Gap-Filling

First, we downloaded the annotated genome sequences of the five external models from NCBI and their GEM SBML format with the reference file of ID reactions present in KEGG, Bigg or MetaCyc. Reconstruction by orthology provided 174 reactions for the *F. succinogenes* S85 first model obtained from Pathway Tools (Fig. 2A). In addition, 203 reactions were brought by the combination of reconstructions by annotation and orthology. Finally, 61% of orthology-based reactions were added to the network according to cross-sources (Fig. 2B).

The definition of seeds and targets is an essential step in the reconstruction protocol. We have determined a list of 51 sources (constituents of the culture medium) and 85 targets (molecules known to be produced by the bacterium) for *F. succinogenes* S85 (Table S2 in Supplementary Material A). The gaps in the model were firstly filled automatically by mapping and transforming KEGG/Bigg identifiers to MetaCyc IDs using the MetaNetX package. 99 reactions were added by Meneco (40), all of them came from the MetaCyc database (45). 24 reactions were removed according to expert validation (see below), finally 75 reactions were added by gap-filling.

#### • FBA for unblocking biomass and manual curation

In our reconstruction process, the aim was to obtain a functional high-quality network, which produces biomass yield. For this purpose, we firstly focused on reaching topologically all the targets according to the qualitative network-expansion criteria. Forty-one reactions were expertly added to the gap-filled model network. This reactions belong to four distinct databases: MetaCyc (45), KEGG(63), BIGG(64) and RHEA (65).

Afterward, we manually unlocked all the pathways referring to FBA analysis, which were in direct and indirect relation with each component of the biomass reaction. We therefore focused on curating the network and the pathways involved in carbon and nitrogen metabolism as well as some essential cofactor biosynthesis. In particular, we checked the different pathways of glycolysis, glycogen cycle (see below) and the short chain fatty acids biosynthesis. This required adding 490 manually curated reactions in the pathways of the biomass compounds.

#### • Manual completion

*F. succinogenes* S85 model reactions are linked to 1317 genes, leading to 77% of reactions associated to a gene (Fig.3). 430 reactions were not linked to any gene before validation. Of them, 133 are exchanged, transport or spontaneous reactions. For all the other reactions, we search for the presence of a gene possibly associated using BLAST (Data set S1 in Supplementary Material B). Finally, 68 of those were added by Gap-filling (Fig. 2A), 54 reactions that seemed inappropriate to the metabolism of the bacterium were suppressed, and 22 reactions were linked manually to their corresponding gene.

**Figure 3.**
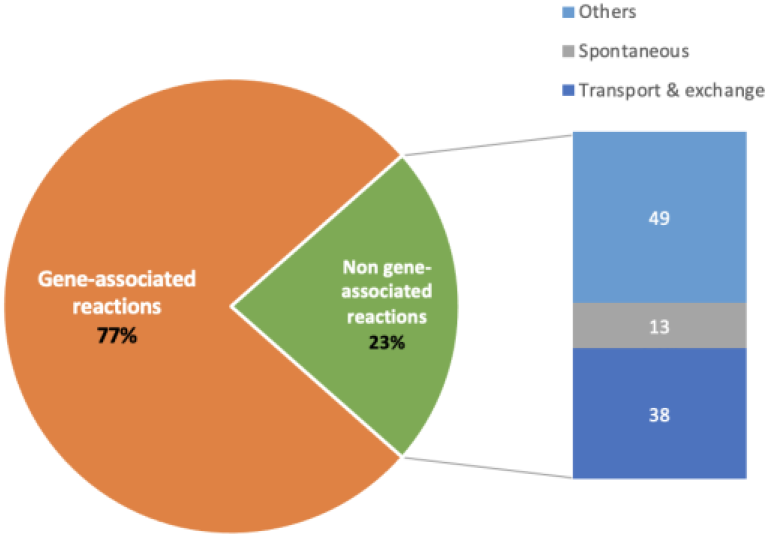
Gene – reactions association.

### 3.2. Qualitative analysis of the *Fibrobacter succinogenes* S85 metabolic network

#### 3.2.1. FBA and essential reactions

The obtained network has 1317 genes, 931 pathways and is composed of 1565 reactions, from which 1211 are associated to genes (Table 1). The final network contains 1586 unique metabolites and is available online as a wiki page on: https://gem-aureme.genouest.org/fsucgem/index.php/Fsucgem.

**Table 1.** *F. succinogenes* S85 metabolic model information.

|  | *F. succinogenes* S85 model |  |  |  |
| --- | --- | --- | --- | --- |
| Reactions | 1565 |  |  |  |
| Unique metabolites | 1586 |  |  |  |
| Genes | 1317 |  |  |  |
| Active* / Total pathways | 233/931 |  |  |  |
|  | Number of reactions** |  |  |  |
|  | Annotation<br>Total: 827 | Orthology<br>Total: 377 | Gap-filling<br>Total: 75 | Manual curation<br>Total: 490 |
| Exchange / Transporter | 1 | 15 | 0 | 133 |
| Spontaneous | 1 | 2 | 1 | 45 |
| Protein / Amino acids biosynthesis | 101 | 74 | 16 | 25 |
| Glycolysis / Fatty acids biosynthesis | 51 | 17 | 2 | 12 |

\* Ratio (Reaction found / Total) > 0,75 and ratio reaction with flux/reaction based on FBA analysis > 0,5
\*\*204 reactions are identified from cross method sources

We investigate our final metabolic network using Flux Variability Analysis (FVA). All the 85 target components are reached topologically. 38,5% of the reactions are active, of which 137 are essential reactions for the biomass production. The simulated growth rate from *F. succinogenes* S85 metabolic model is 0.137 h^-1^. 318 out of 931 (34%) of the metabolic pathways are complete at more than 75% of the reactions present in the KEGG and MetaCyc databases. 76% of those active pathways have at least one reaction with flux according to the FBA analysis, and 150 pathways are 100% active in flow (Data set S2 in Supplementary Material B). All the essential reactions are present in the active pathways.

#### 3.2.2 Glycogen biosynthesis and degradation pathways

As an example to illustrate the use of the reconstructed network, we analyzed the glycogen biosynthesis and degradation pathways, because glycogen has been shown to be simultaneously synthesized and degraded in *F. succinogenes* during all growth phases (19). Intracellular glycogen accumulation is carried out by the consecutive action of ADP-glucose pyrophosphorylase (*glgC*, FISUC_RS14455 FSU_RS00645) (EC.2.7.7.27), glycogen synthase *(glgA* FSU_RS16140 FISUC_RS15965) (EC.2.4.1.21), and glycogen branching enzyme *(glgB* FISUC_RS15575 FSU_RS01770) (EC.2.4.1.18) (Fig. 4). All these anabolism reactions and genes were identified by annotation, except the phosphoglucomutase (*pgm* FSU_0773) that was linked to the reaction EC 5.4.2.2 by manual curation (Data set S1 in Supplementary Material B).

**Figure 4.**
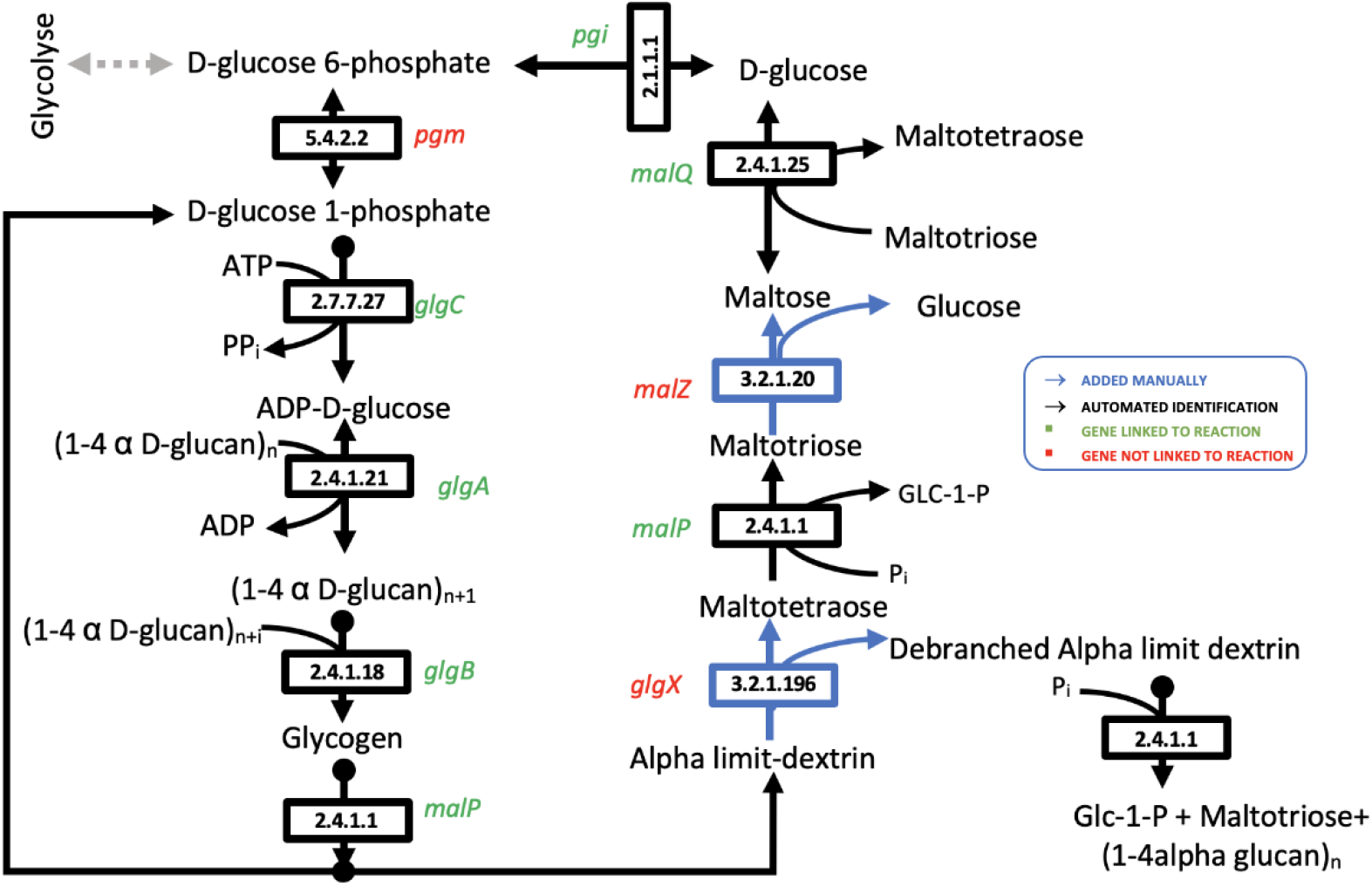
Futile glycogen pathways identified and completed in F. succinogenes S85 metabolic network.

The known glycogen degradation I pathway on MetaCyc 23.0 was detected with the presence of the reactions EC 2.4.1.1 (Maltotetraose glucosidase *malP:* FSU_RS06195) and EC 2.4.1.25 (4-alpha-glucanotransferase *(malQ* FISUC_RS04305; FSU_RS06200) identified by annotation (Fig. 4). Then, we completed this pathway by adding manually the two reactions EC 3.2.1.196 (limit dextrin α-1,6-glucohydrolase *glgX)* and EC 3.2.1.20 alpha-glucosidase *(malZ* FSU_RS06195). The maltotetraose formation reaction present in *Escherichia coli* model was added to our model, no gene was linked because of no significant blast similarity with the *F. succinogenes* genome (Data set S1 in Supplementary Material B).

### 3.3. Network reduction

#### ■ Reduced-scale genome reconstruction model process

Network reduction was achieved by NetRed method which is based on flux vectors computed by FBA from the full network. The reactions in the full network were defined as irreversible reactions by decoupling the reversible reaction into their forward and backwards directions. The full network was then composed by 1780 irreversible reactions. The measured metabolites obtained from batch cultures growing on glucose, cellobiose and cellulose corresponded to: extracellular acetate, formate and succinate. From those results, it was observed that formate is produced in small amounts while succinate and acetate were the main products. Accordingly, two objective functions were defined to maximize concomitantly: (1) biomass and succinate; and (2) biomass and acetate. In order to enlarge the possible flux distribution, different input fluxes of glucose, cellobiose and cellulose ranging between 0 to 1000 mmol/g_biomass_ h^-1^ with a step of 50 mmol/g_biomass_ h^-1^ were considered. For the sake of computation ease, we defined that cellobiose was composed of 2 glucoses while cellulose was considered as 4 glucoses, so the upper ranges were modified to 250 and 500 mmol/g_biomass_ h^-1^ respectively. The resultant fluxes were analyzed in terms of yields for which all fluxes were divided by the flux of the carbon uptake reactions.

Carbon balance was verified through yield analysis, where we have noticed that the glycogen synthesis and degradation pathways generated unbalanced carbon production. For the sake of carbon quantification, we assumed that glycogen and (1,4-α-D-glucan)_n_ were composed of 6 and 5 molecules of glucose, respectively. Additionally, a new hypothetical reaction was added to consider the production of (1,4-α-D-glucan)_n_ as,

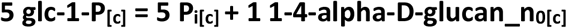

This new reaction allowed to balance carbon and produce biomass with a mass of 25.394 g/mol_c_. FBA was performed again to compute the flux vector and performed the reduction via NetRed. The protected metabolites corresponded to: biomass, acetate, succinate, formate, glycogen, glucose, cellulose, cellobiose, protons, ammonium and fructose-6P. The reduced network contained 146 reaction and 78 metabolites whose size was still large for computation of EFM.

A further reduction of the network was achieved by constructing a lumped biomass reaction which was based on the precursors of several metabolites such as amino acids. The precursors were identified by tracking back the pathways that produce the metabolite to be deleted from the 137 essential reactions identified for the full network. All the reactions involved in the pathways were added up to obtain a reaction to replace the metabolite to be deleted by its precursors (e.g., pyruvate, fructose-6P). This approach is similar to the computing of lumped biomass employed by Lugar et al., (56). Details about the construction of the lumped biomass could be found in the Supplementary Material B (Data set S3 in Supplementary Material B). Once the lumped biomass reaction was obtained, the coefficients were corrected to obtain a biomass of 26.401 g/mol_c_ following the formula C_3.69_H_6.76_O_2.66_N_0.25_S_0.010_ previously reported for the biomass composition of *F. succinogenes* S85 (66).

The new network comprising the lumped biomass reaction and the carbon balance was used to calculate flux vectors subject to several constraints. All the reactions used for the construction of the lumped biomass reaction that did not correspond to essential reaction were blocked to zero flux. Furthermore, we verified that cofactors such as FAD, P_i_, PP_i_, and ADP were not needed as sources, so that their fluxes were also assumed to be zero. A reduction of the extracellular cofactors was made accepting small changes in yield analysis from FBA. Cofactors such as NADPH, NADP, NAD, NADH were not needed as sources, whereas ADP and NADH were not needed as sinks.

Flux vectors for the three substrates and the protected metabolites mentioned before were used in NetRed to obtain a reduced network of 63 reactions with 36 intracellular metabolites and 16 extracellular metabolites. The biomass reaction of the reduced network accounts for a term called ‘salts’ gathering all the metabolites that do not participate in any other reactions, but that are, nevertheless, necessary to produce biomass. The extracellular metabolites were biomass, acetate, succinate, formate, ammonium, Co-A, CO_2_, proton (cytosolic and external), ATP, salts, glycogen, PP_i_ and the three carbon sources glucose, cellobiose and cellulose. The reduced network (Data set S4 in Supplementary Material B) is appropriate for the computation of EFM.

### 3.4. From EFMs to macroscopic reactions

EFMs were computed for each carbon source obtaining 9 861 037, 11 863 589 and 11 540 721 EFMs for glucose, cellobiose and cellulose, respectively. The calculation did not consider that the network could use the three carbon sources at the same time. For the sake of analysis, only the EFMs that consumed the carbon source, produced biomass, and did not consume glycogen were considered for analysis leading to a total of 798 872, 1 198 271 and 2 131 696 EFMs for glucose, cellobiose and cellulose, respectively.

The computed EFMs were multiplied by the stochiometric matrix of the extracellular metabolites to derive macroscopic reactions that can efficiently bring together metabolism and dynamics through the development of Dynamic Metabolic Models (DMM) (67, 68). However, the consideration of many EFMs adds considerably to the kinetic parameters associated with substrate uptake rates leading to an overparameterization. Yield analysis (67) was presented as an alternative to perform a substantial reduction of the number of EFM from an inspection of the convex hull in a 2-D representation on the yield vector space for extracellular products.

Yield analysis for the EFMs is reported for the four principal products: biomass, acetate, formate, and succinate. Yields of the computed EFMs were obtained by dividing their fluxes by the flux of the carbon source. The minima and maxima yield values obtained from EFM for the main products are reported in Table 2.

**Table 2.** Yield boundaries of the reduced Network.

|  | Yield | Biomass | Acetate | Succinate | Formate |
| --- | --- | --- | --- | --- | --- |
| **Glucose** | **min** | 0.0002 | 0.0000 | 0.0000 | 0.0000 |
|  | **max** | 0.1465 | 2.6231 | 1.5622 | 3.8813 |
| **Cellobiose** | **min** | 0.0001 | 0.0000 | 0.0000 | 0.0000 |
|  | **max** | 0.2935 | 5.1987 | 3.1664 | 4.5788 |
| **Cellulose** | **min** | 0.0012 | 0.0000 | 0.0000 | 0.0000 |
|  | **max** | 0.5801 | 6.6722 | 6.2504 | 4.9125 |

Fig. 5 displays in the diagonal the distribution of the 798 872 EFM obtained for glucose where it is observed that biomass is mainly produced at values around 0.01 g per mmol of glucose. On the other hand, most of the EFMs producing acetate, succinate, formate report small extracellular production. The plots in the non-diagonal show the yields of the products with respect to all the products where each blue point is an EFM. It is worth noting that the surfaces in yields mainly correspond to triangles except for the yields for formate. Similar results were obtained when the only carbon source was cellobiose (Fig. 6) and cellulose (Fig. 7).

**Figure 5.**
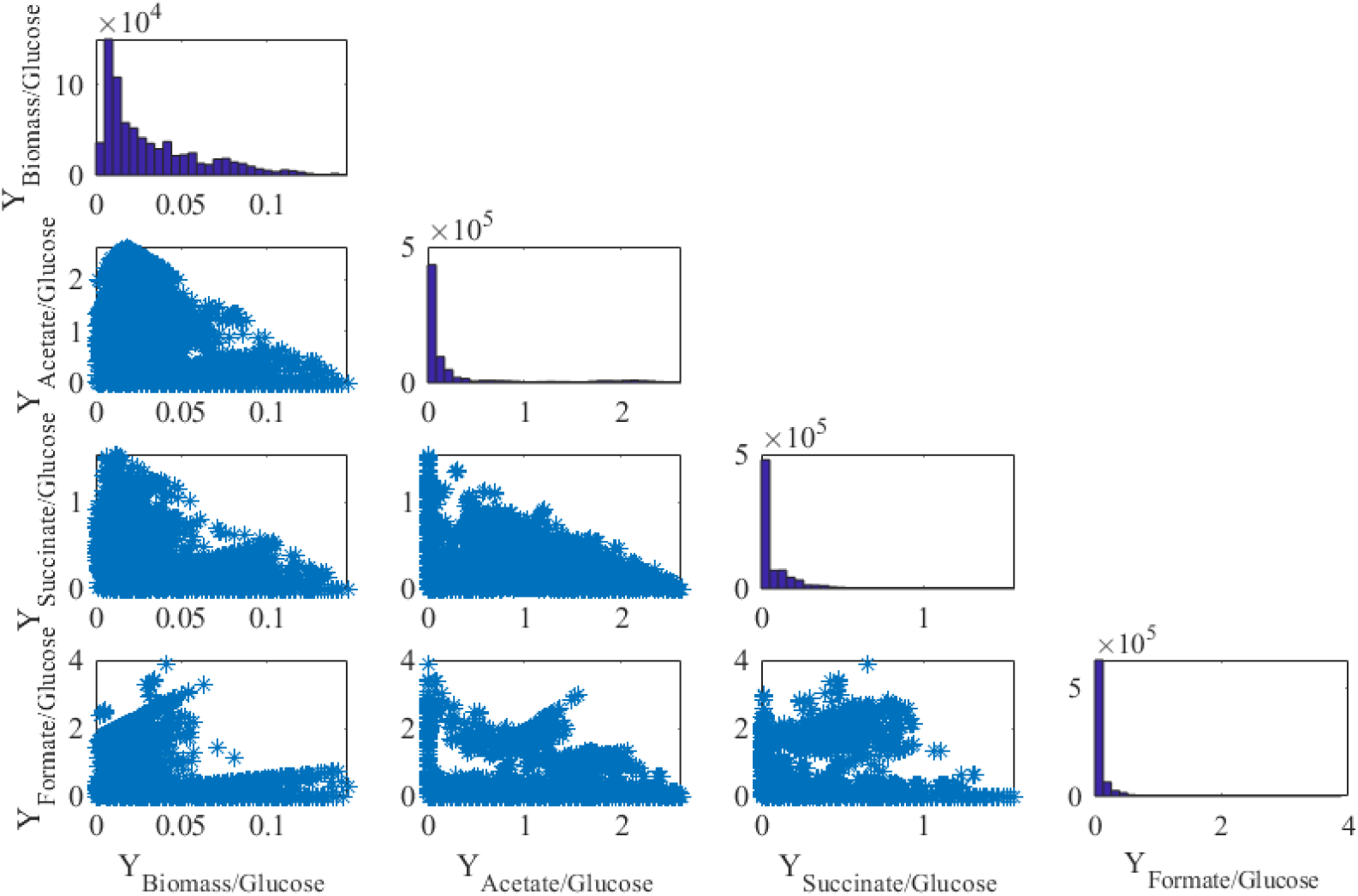
Yield representation of EFM and distribution of EFM when growing on glucose. Units are mmol/mol except for the yield biomass/glucose (g/mmol).

**Figure 6.**
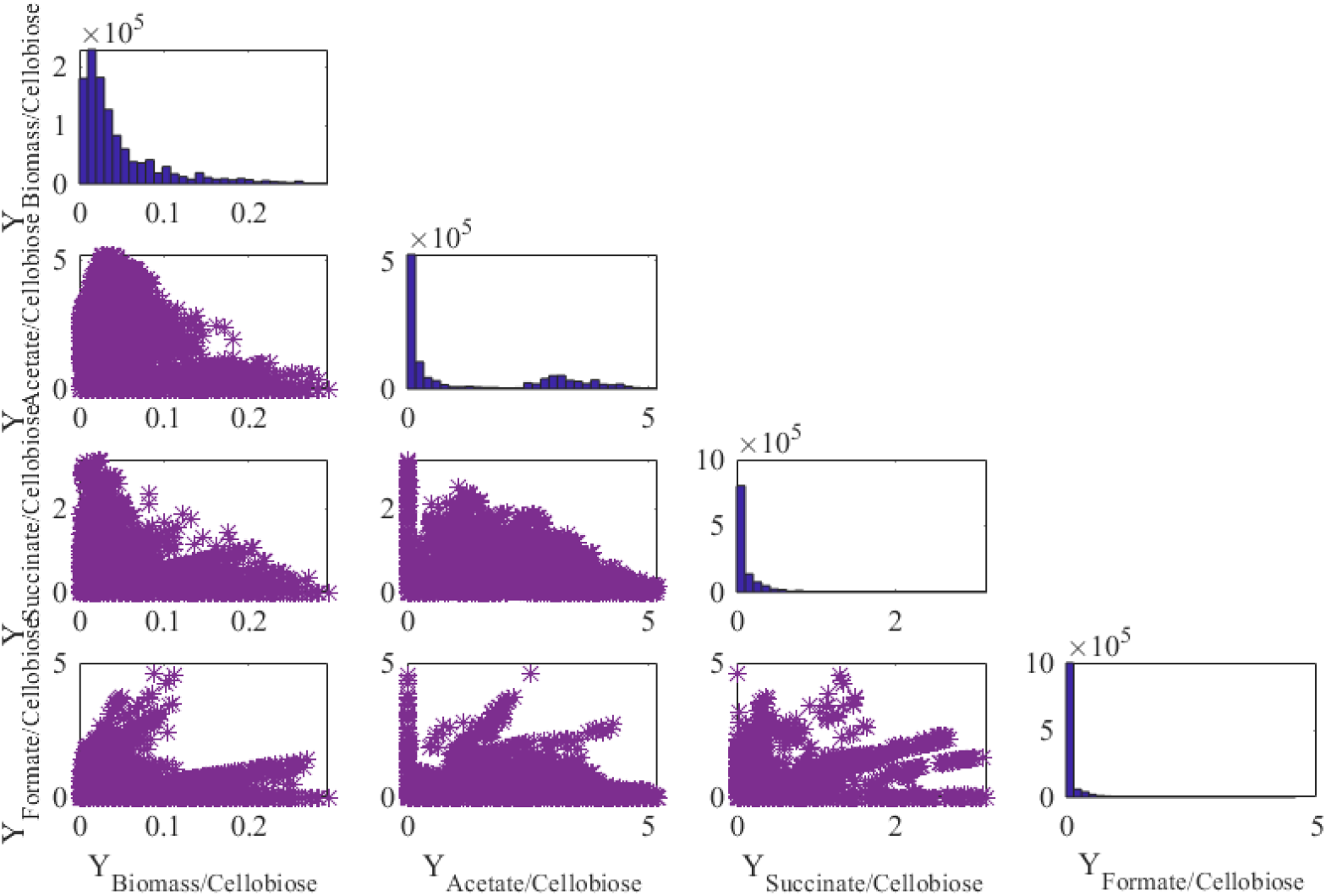
Yield representation and distribution of EFM when growing on cellobiose. Units are mmol/mol except for the yield biomass/cellobiose (g/mmol).

**Figure 7.**
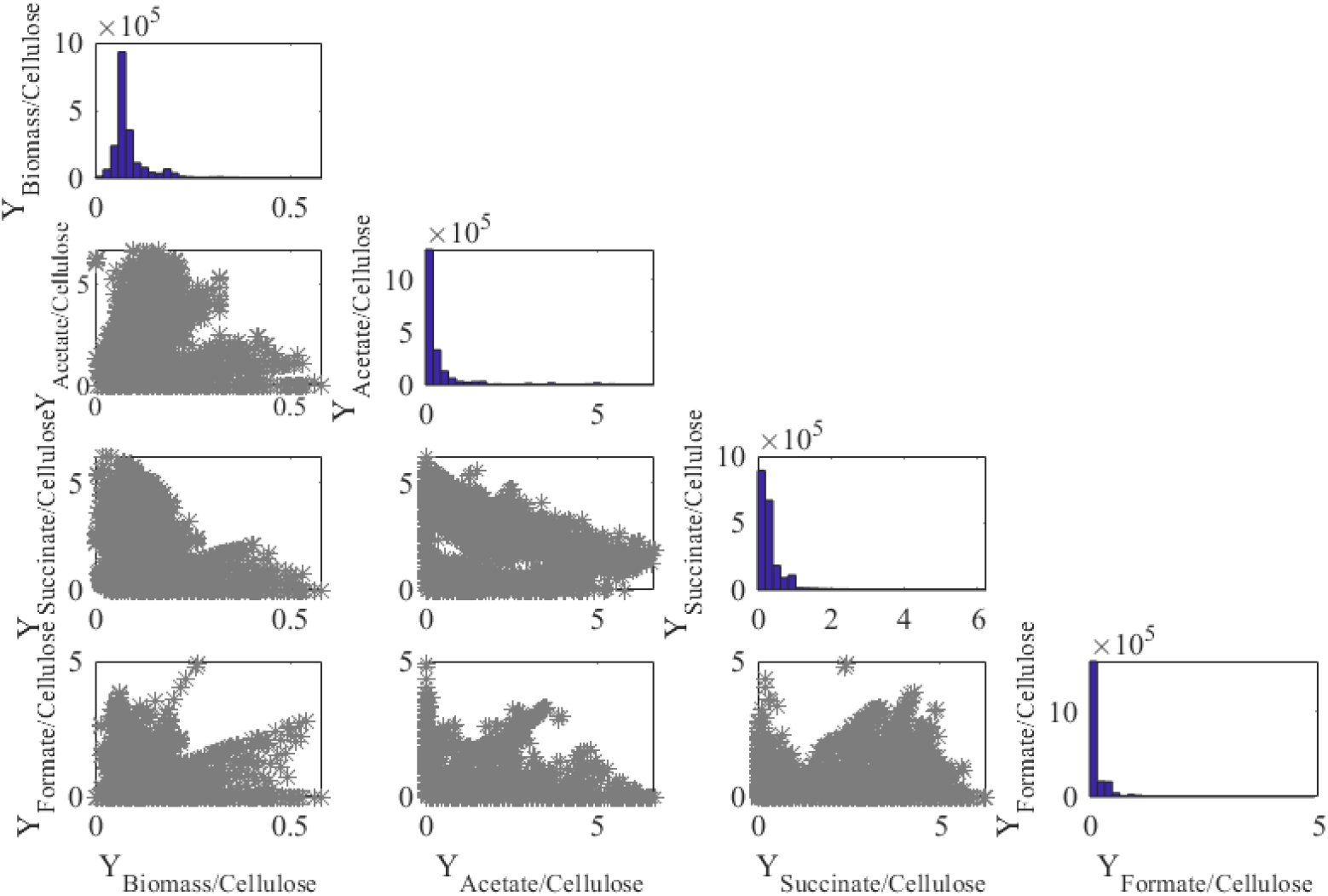
Yield representation and distribution of EFM when growing on cellulose. Units are mmol/mol except for the yield biomass/cellulose (g/mmol).

We performed a yield analysis with a 2D representation of the convex hull -surrounding total EFMs- to reduce the number of EFM used for macroscopic reactions of DMM. In this case, triangles were used to find a minimum number of EFM to be used in DMM. Those EFM are denoted as red points in Figures 8 A-C which display the EFM and their reduction by yield analysis for glucose, cellobiose and cellulose, respectively.

**Figure 8.**
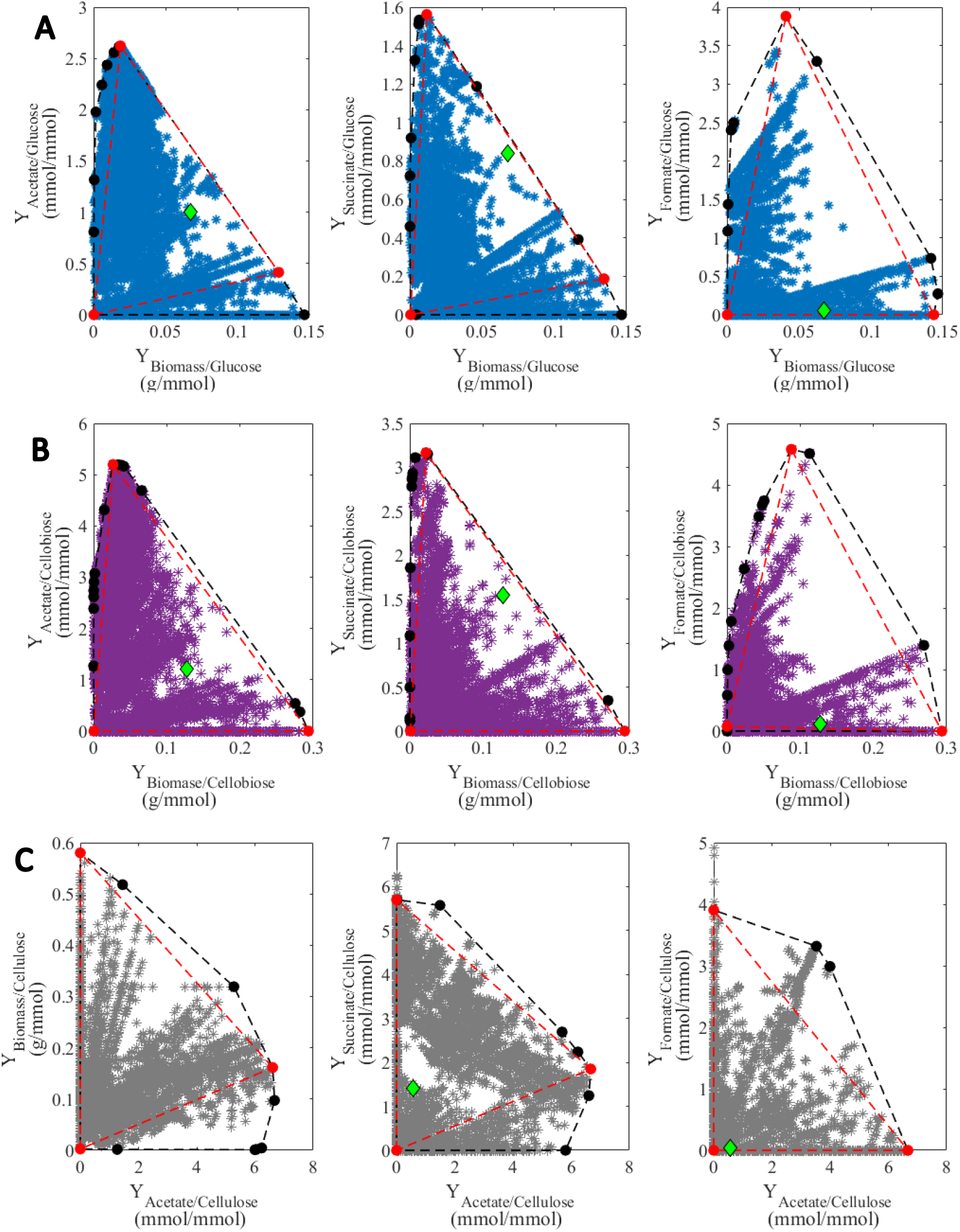
Yield analysis of the EFMfor A 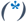 glucose B 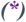 cellobiose and C 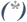 cellulose. Representation computation of the convex hull (-•-) and reduction of the convex hull 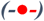 with respect to experimental data 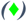.

The 9 selected EFM obtained for each substrate were compared to avoid repeated EFM. For glucose, 9 EFM remained while only 7 and 8 remained for cellobiose and cellulose (Table 3).

**Table 3.**
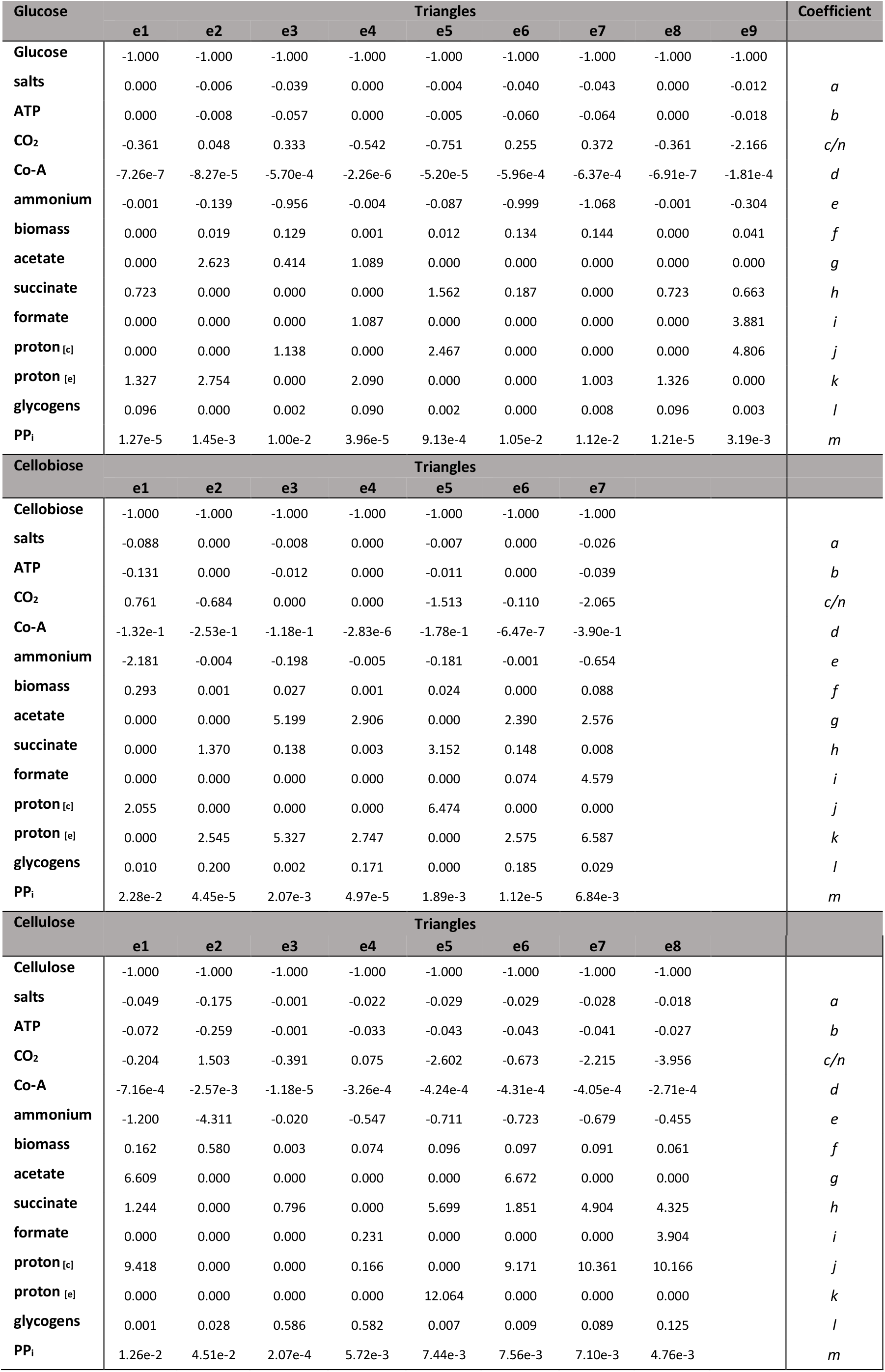
EFMs from the polygon (triangle) enclosing the yield spaces.

From Table 3, macroscopic reactions for each substrate can be derived as:

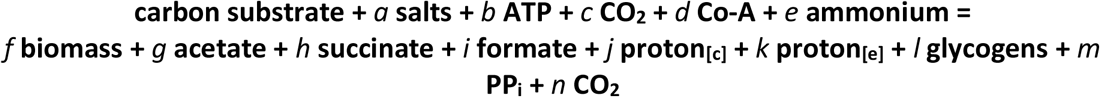

where the coefficients *a – n* correspond to the absolute values of the EFM which represent the letters on the Table. Note that the coefficients *c* or *n* will depend on whether CO_2_ has a negative sign (*c*) or positive sign (*n*). These EFMs are used to select a minimal set of macroscopic reactions for the dynamic metabolic model (DMM) as discussed below.

### 3.5. Dynamic metabolic model

Table 4 shows the selected EFMs, and the model parameter estimates for each substrate. The metabolism of glucose and cellulose is represented by 4 macroscopic reactions. For cellobiose, the metabolism is represented by 5 macroscopic reactions. All the EFMs are from the polygon vertices of the yield spaces. The models were implemented in MATLAB and are available at https://doi.org/10.5281/zenodo.7219865.

**Table 4.**
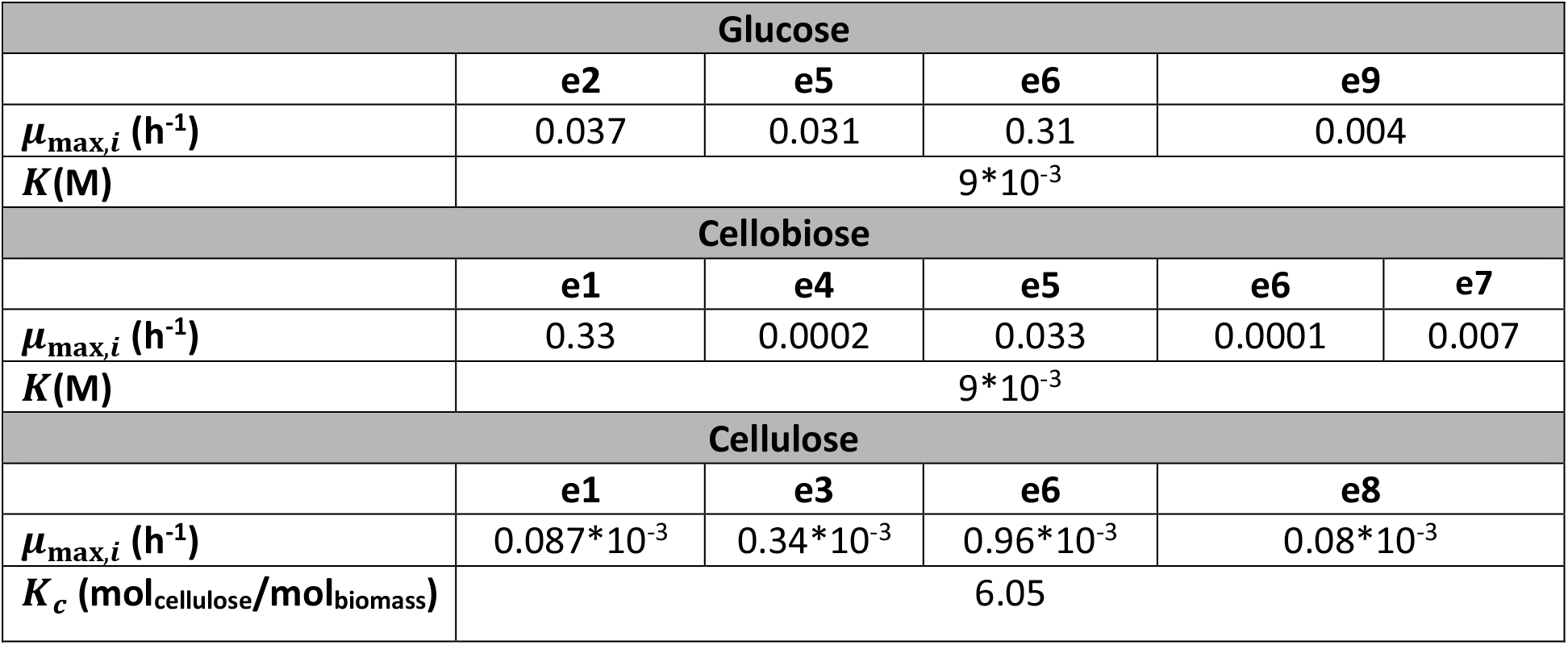
Selected EFMs of the dynamic model and parameters estimates. The stoichiometry of the EFMs is given in Table 3.

| Glucose |  |  |  |  |  |
| --- | --- | --- | --- | --- | --- |
|  | e2 | e5 | e6 | e9 |  |
| $\mu_{\max,i}$ (h<sup>-1</sup>) | 0.037 | 0.031 | 0.31 | 0.004 | |
| $K(\text{M})$ | 9*10<sup>-3</sup> | | | | |
| Cellobiose |  |  |  |  |  |
|  | e1 | e4 | e5 | e6 | e7 |
| $\mu_{\max,i}$ (h<sup>-1</sup>) | 0.33 | 0.0002 | 0.033 | 0.0001 | 0.007 |
| $K(\text{M})$ | 9*10<sup>-3</sup> | | | | |
| Cellulose |  |  |  |  |  |
|  | e1 | e3 | e6 | e8 |  |
| $\mu_{\max,i}$ (h<sup>-1</sup>) | 0.087*10<sup>-3</sup> | 0.34*10<sup>-3</sup> | 0.96*10<sup>-3</sup> | 0.08*10<sup>-3</sup> | |
| $K_c$ (mol<sub>cellulose</sub>/mol<sub>biomass</sub>) | 6.05 | | | | |

Figures 9A-C show the comparison of the experimental data against the variables predicted by the model. Table 5 shows the accuracy of the model. For the experiments with glucose, the coefficient of variation of the Root mean squared error CVRMSE was 17%. For the experiments with cellobiose, the average CVRMSE was 19%. For the experiments with cellulose, the average CVRMSE was 22%.

**Figure 9.**
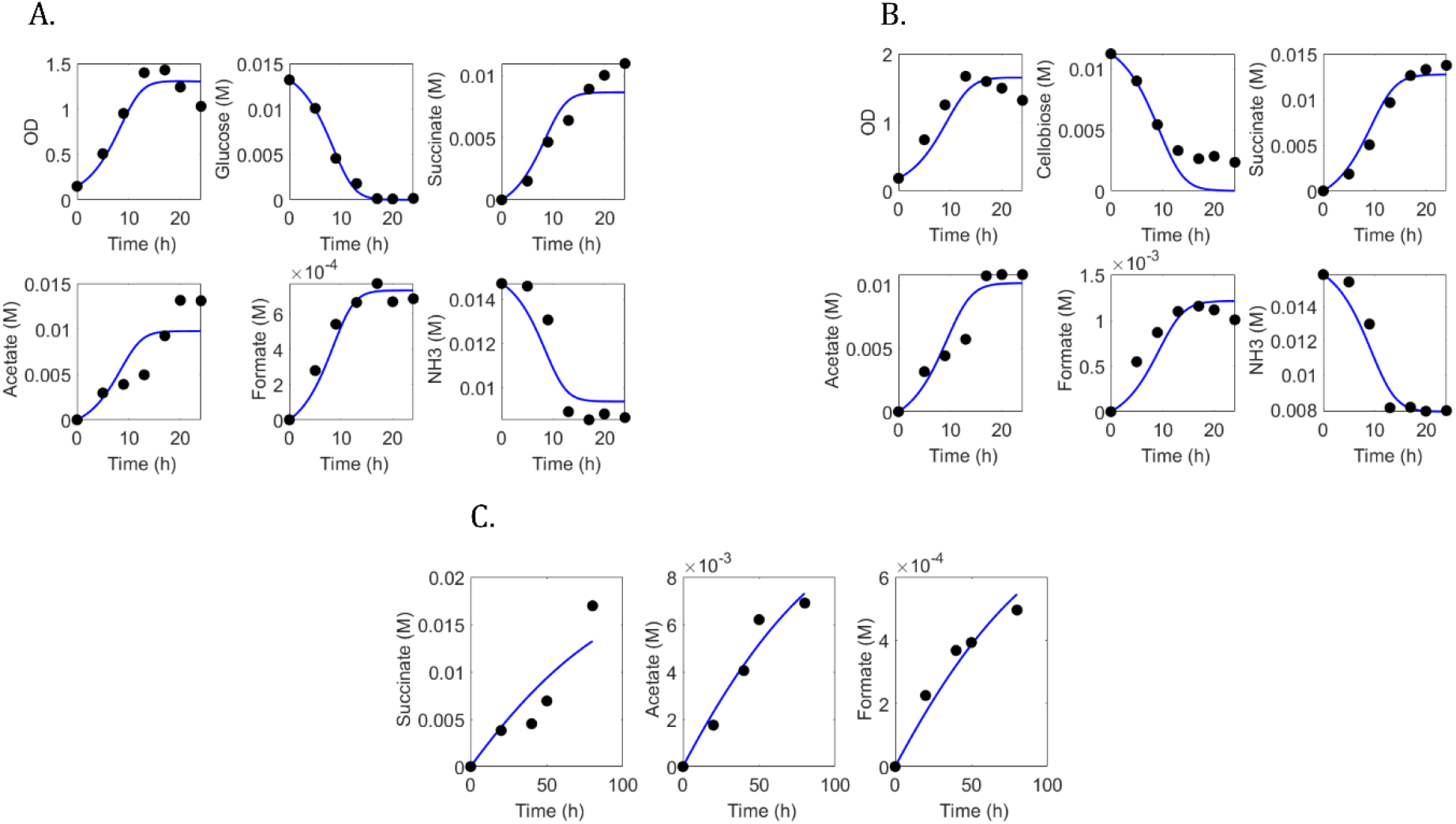
Experimental data (•) of the fermentation of (A) glucose (B) cellobiose (C) cellulose compared against the variables predicted by the dynamic models (solid line).

**Table 5.** Model accuracy.

|  | Glucose utilization |  |  |  |  |  |
| --- | --- | --- | --- | --- | --- | --- |
|  | Acetate | Succinate | Formate | Ammonia | Substrate | OD |
| 100*CVRMSE<sup>a</sup> | 34 | 19 | 11 | 10 | 14 | 14 |
|  | Cellobiose utilization |  |  |  |  |  |
|  | Acetate | Succinate | Formate | Ammonia | Substrate | OD |
| 100*CVRMSE<sup>a</sup> | 18 | 11 | 19 | 8 | 44 | 17 |
|  | Cellulose utilization |  |  |  |  |  |
|  | Acetate |  | Succinate |  | Formate |  |
| 100*CVRMSE<sup>a</sup> | 15 |  | 38 |  | 13 |  |

<sup>a</sup> Coefficient of variation of the RMSE (CV(RMSE)).

## 4. Discussion

The mathematical modelling of the rumen ecosystem is a useful endeavor to provide tools for improving rumen function. Current kinetic rumen models do not consider genomic information (31, 59, 69, 70). GEMs are a promising tool to fill this lacking gap and allowing a better understanding of the rumen systemic functionality (71) and the individual bacteria metabolism (72).

Many independent methods have been developed to generate genome scale models, including some toolboxes and workspaces, such as Pathway Tools (36), RAVEN (73), merlin (74), KBase (75), The SEED (76), AuReMe (35), AutoKEGGRec (63), CarVeMe (77), and gapseq (78). They rely on one or several metabolic databases such as MetaCyc(45), KEGG(79), ModelSEED(80) or BiGG (64). However, the output of a main platform for a GEM requires adjustments assisted by a choice of specialized tools, especially when the network reconstruction requires to take advantages of information spread in different models, formats, and organisms, leading to issues in standardization of metadata and reproducibility of the reconstruction procedure. In this work, the GEM construction of *Fibrobacter succinogenes* S85 was performed using the AuReMe platform, selected for its full traceability reconstruction (35) and capabilities to produce high-quality reconstructions (81).

Genome-scale metabolic models are widely used for microbial defined growth medium identification (30, 82), metabolic functional characterization (29, 83) or design of novel treatment against pathogens (84). Regarding gut communities, they have been mainly applied to human gut bacteria in order to decipher the microbial interactions in the human intestinal microbiome (28, 30, 85–87).

Until now there have been few GEMs available for rumen bacteria such as the networks for the lactate utilizing bacterium *Megaspghera elsdenii* (72), and the succinic acid producing strain *Actinobacillus succinogenes* (88). Recently, one simplified representative rumen community metabolic model was reported (71). A synthetic community composed of a cellulolytic bacterium, a proteolytic bacterium and a methanogen was developed to enlighten metabolite secretion profiles, community compositions and the interactions with bacteriophages (71). Our work contributes to expand the application of the GEM approach to study the rumen ecosystem.

The obtained GEM of *F. succinogenes* is composed of thousands of metabolites and reactions associated to their genes and can set a useful network information for generating future ruminal bacterial draft models. The 1317 genes of the final *F. succinogenes* S85 model cover more than 41.5% of the genes identified in the two genomes of the strain (3170 and 3161 ASM14650v1 and ASM2466v1, respectively).

Our model contains 2.5 times more genes than the model of the rumen cellulolytic bacterium *Ruminococcus flavefaciens* previously reconstructed (71) by ModelSEED (80) and gaps-filled by GapFind-GapFill (71), as well as 1.5 more reactions and metabolites. Our final reconstructed network is functional for biomass and SCFA production with compacted core metabolism represented by 30% of active pathways (Data set S3 in Supplementary Material B) and has a simulated growth rates 72% times greater than that of *R. flavefaciens*, its cellulolytic model candidate (71). GEMs are very powerful to provide a qualitative analysis of microbial metabolism. However, they are limited for quantitative prediction of the dynamics of metabolites. This work develops an approach for developing a dynamic metabolic model (DMM) exploiting the microbial genomic information embedded in the GEM of *F. succinogenes* S85. The use of GEM for dynamic modeling and other analysis methods is cumbersome due to the large number of reactions and metabolites which hampers the interpretation/visualization of fluxes (e.g., FBA) and limits the calculation of computationally expensive analysis (e.g. Elementary Flux Modes (EFM)) (89). Hence, a reduction of the GEM into a network that still captures phenotypic and genotypic properties while displaying flexibility is needed (90). For our modelling exercise, the NetRed tool (56) was instrumental to perform the reduction of our network. All the full-scale GEM reactions participating in the 63 reduced-scale genome based metabolic network reactions of *F. succinogenes* S85 are present in the list of active pathways of the large-scale network (Data set S2 in Supplementary Material B).

Yield analysis was used in the flux vectors to verify the carbon balance of the network. The obtained yields showed that carbon balance was not respected, and the difference was coming from the glycogen pathways. Some of the glycogen pathway reactions were identified from the databases as generic reactions. *F. succinogenes* is known to synthesize glycogen during all its growth phases (19). We focused on the validation of its production/degradation pathway including generic reactions not only for its specificity in this organism but also for its importance in the carbon cycle equilibrium that needs to be held for the network, the EFM computation step and the dynamic metabolic model. As solution, we have set for glycogen a number of monomers equal to 6 in order to be able to complete the carbon balance and therefore stoichiometrically balance the reactions of this metabolic pathway. The reduced network helped to calculate the EFM whose reduction was performed by yield analysis (67). A 2-D representation of yields was used to compute the convex hull that surround the EFMs. The EFMs belonging to the convex hull were further reduced by a method employing polygons (91). The EFMs on the convex hull are normally chosen to provide a wide range of steady states to the system.

The resulting model structures are similar in degree of complexity with respect to the rumen fermentation model developed by Muñoz-Tamayo et al (92) where carbohydrate metabolism is represented by 5 macroscopic reactions. The main difference in the approach developed in the present work is that the macroscopic reactions are derived from the reconstructed metabolic network of *F. succinogenes*. It should be noted that the resulting macroscopic reactions included in the models correspond to the active sets of EFMs specific to the experimental here studied while the subsets of all EFMs at the vertices of the polygon enclosing the yield spaces shown in figures 8A-C constitute a minimal generating of EFMs covering almost all possible metabolic states. This approach provides a high flexibility to span the metabolic space at different experimental conditions. Such a flexibility in the model structure is a great asset to study in the future strategies to enhance substrate utilization and target desired fermentation profile.

The model performances were acceptable to capture the dynamics of fermentation by *F. succinogenes*. However, Table 5 and Figures 9A-C display that there is room for improvement. The prediction of succinate by the model for cellulose utilization has indeed a high CVRMSE. One key element for model improvement is glycogen metabolism, which was not integrated in this work. Glycogen plays an important role in *F. succinogenes* and appears to be submitted to a futile cycle which results from a simultaneous utilisation and storage (19, 93). As observed in Table 3, glycogen is a net product for the EFMs of the polygon vertices. Thereby, the current model structure cannot account for the futile cycling. This limitation is intrinsic to the steady state assumption for the EFM derivation. To account for glycogen futile cycling, it will be then required to split the metabolic network into subnetworks. The procedure of network splitting can be done on knowledge basis as applied for example to study microalgae metabolism (94). However, the splitting method is a challenging issue. As perspective, in the mid-term, we will explore the use of splitting techniques such as those developed by (95, 96) to account for the glycogen futile cycle. In the long-term, we will apply the approach here developed to other key rumen microbes to address the modelling of rumen microbial mini consortia. As we have previously discussed (97), this approach will enable us to construct tractable models that integrate genomic information with capabilities to inform on strategies for driving the rumen microbiome.

## 5. Acknowledgements

We thank the INRAE PHASE department, and the INRAE MEM metaprogram for financial support. The PhD of Ibrahim Fakih is supported by grants from the INRAE Holoflux metaprogram and funding from Lallemand Animal Nutrition (Blagnac, France).

We thank Emile Dumont, Méziane Aite for their preliminary work on network reconstruction, and Melanie Brunel for the cultivation experiments with *F. succinogenes*.

## Availability of data and material

The metabolic networks and mathematical models developed in this work are freely available at https://doi.org/10.5281/zenodo.7219865.

### Supplementary material A

□ Table S1 Biomass composition
□ Table S2 List of seeds and targets
□ Table S3 Comparison between the two draft models obtained via annotation

### Supplementary material B

□ Data set S1 Blast results for gene-reaction association
□ Data set S2 List of active pathways
□ Data set S3 List of spontaneous, transport reactions
□ Data set S4 Lumped biomass reaction
□ Data set S5 Reduced model

## **Authors’ contributions (**Using the taxonomy of CREDIT)

**Ibrahim Fakih:** Conceptualization, Investigation, Methodology, Writing – original draft

**Jeanne Got:** Conceptualization, Investigation, Methodology, Software, Writing – review & editing

**Carlos Eduardo Robles-Rodriguez:** Conceptualization, Investigation, Methodology, Software, Writing – review & editing

**Anne Siegel:** Conceptualization, Methodology, Writing – review & editing, Supervision

**Evelyne Forano:** Conceptualization, Methodology, Writing – review & editing, Supervision, Funding acquisition,

**Rafael Muñoz-Tamayo:** Conceptualization, Methodology, Software, Writing – review & editing, Supervision, Funding acquisition, Project administration

## References

1. Huws SA, Creevey CJ, Oyama LB, Mizrahi I, Denman SE, Popova M, Muñoz-Tamayo R, Forano E, Waters SM, Hess M, Tapio I, Smidt H, Krizsan SJ, Yáñez-Ruiz DR, Belanche A, Guan L, Gruninger RJ, McAllister TA, Newbold CJ, Roehe R, Dewhurst RJ, Snelling TJ, Watson M, Suen G, Hart EH, Kingston-Smith AH, Scollan ND, Prado RMD, Pilau EJ, Mantovani HC, Attwood GT, Edwards JE, McEwan NR, Morrisson S, Mayorga OL, Elliott C, Morgavi DP. 2018. Addressing global ruminant agricultural challenges through understanding the rumen microbiome: Past, present, and future. Frontiers in Microbiology 9:1–33.

2. Moraïs S, Mizrahi I. 2019. The road not taken: The rumen microbiome, functional groups, and community states. Trends in Microbiology 27:538–549.

3. Xie S, Syrenne R, Sun S, Yuan JS. 2014. Exploration of Natural Biomass Utilization Systems (NBUS) for advanced biofuel–from systems biology to synthetic design. Current opinion in biotechnology 27:195–203.

4. Weimer PJ. 1996. Why don’t ruminal bacteria digest cellulose faster? Journal of Dairy Science 79:1496–1502.

5. Henderson G, Cox F, Ganesh S, Jonker A, Young W, Janssen PH, Collaborators GRC. 2015. Rumen microbial community composition varies with diet and host, but a core microbiome is found across a wide geographical range. Sci Rep 5:14567.

6. Neumann AP, McCormick CA, Suen G. 2017. *Fibrobacter* communities in the gastrointestinal tracts of diverse hindgut-fermenting herbivores are distinct from those of the rumen. Environmental microbiology, 2017/08/24 ed. 19:3768–3783.

7. Hungate RE. 1950. The anaerobic mesophilic cellulolytic bacteria. Bacteriological reviews 14:1–49.

8. Bryant MP, Doetsch RN. 1954. A study of actively cellulolytic rod-shaped bacteria of the bovine rumen. Journal of Dairy Science 37:1176–1183.

9. Raut MP, Couto N, Karunakaran E, Biggs CA, Wright PC. 2019. Deciphering the unique cellulose degradation mechanism of the ruminal bacterium *Fibrobacter succinogenes* S85. Scientific reports 9:16542.

10. Suen G, Weimer PJ, Stevenson DM, Aylward FO, Boyum J, Deneke J, Drinkwater C, Ivanova NN, Mikhailova N, Chertkov O, Goodwin LA, Currie CR, Mead D, Brumm PJ. 2011. The complete genome sequence of *Fibrobacter succinogenes* s85 reveals a cellulolytic and metabolic specialist. PLoS ONE 6.

11. Yeoman CJ, Fields CJ, Lepercq P, Ruiz P, Forano E, White BA, Mosoni P. 2021. In Vivo Competitions between *Fibrobacter succinogenes*, Ruminococcus flavef*aciens*, and *Ruminoccus albus* in a Gnotobiotic Sheep Model Revealed by Multi-Omic Analyses. mBio 12:e03533–20.

12. Matulova M, Nouaille R, Capek P, Péan M, Forano E, Delort AM. 2005. Degradation of wheat straw by *Fibrobacter succinogenes* S85: A liquid- and solid-state nuclear magnetic resonance study. Applied and Environmental Microbiology 71:1247–1253.

13. Shi Y, Weimer PJ. 1996. Utilization of individual cellodextrins by three predominant ruminal cellulolytic bacteria. Applied and environmental microbiology 62:1084–1088.

14. Wells JE, Russell JB, Shi Y, Weimer PJ. 1995. Cellodextrin efflux by the cellulolytic ruminal bacterium *Fibrobacter succinogenes* and its potential role in the growth of nonadherent bacteria. Applied and Environmental Microbiology 61:1757–1762.

15. Béra-Maillet C, Ribot Y, Forano E. 2004. Fiber-degrading systems of different strains of the genus *Fibrobacter*. Applied and Environmental Microbiology 70:2172 LP – 2179.

16. Matte A, Forsberg CW, Gibbins AMV. 1992. Enzymes associated with metabolism of xylose and other pentoses by *Prevotella (Bacteroides) ruminicola* strains, *Selenomonas ruminantium* D, and *Fibrobacter succinogenes* S85. Canadian Journal of Microbiology 38:370–376.

17. Stewart CS, Flint HJ. 1989. Bacteroides (Fibrobacter) succinogen*es*, a cellulolytic anaerobic bacterium from the gastrointestinal tract. Applied Microbiology and Biotechnology 30:433–439.

18. Wells JE, Russell JB. 1994. The endogenous metabolism of *Fibrobacter succinogenes* and its relationship to cellobiose transport, viability and cellulose digestion. Applied Microbiology and Biotechnology 41:471–476.

19. Gaudet G, Forano E, Dauphin G, Delort A-M -M. 1992. Futile cycling of glycogen in *Fibrobacter succinogenes* as shown by in situ1H-NMR and 13C-NMR investigation. European Journal of Biochemistry 207:155–162.

20. Matheron C, Delort AM, Gaudet G, Liptaj T, Forano E. 1999. Interactions between carbon and nitrogen metabolism in *Fibrobacter succinogenes* S85: A 1H and 13C nuclear magnetic resonance and enzymatic study. Applied and Environmental Microbiology 65:1941–1948.

21. Chen B-Y, Wang H-T. 2008. Utility of enzymes from *Fibrobacter succinogenes* and *Prevotella ruminicola* as detergent additives. Journal of Industrial Microbiology and Biotechnology 35:923–930.

22. Arntzen MØ, Várnai A, Mackie RI, Eijsink VGH, Pope PB. 2017. Outer membrane vesicles from *Fibrobacter succinogenes* S85 contain an array of carbohydrate-active enzymes with versatile polysaccharide-degrading capacity. Environmental microbiology 19:2701–2714.

23. Li Q, Siles JA, Thompson IP. 2010. Succinic acid production from orange peel and wheat straw by batch fermentations of *Fibrobacter succinogenes* S85. Applied microbiology and biotechnology 88:671–678.

24. Pais C, Franco-Duarte R, Sampaio P, Wildner J, Carolas A, Figueira D, Ferreira BS. 2016. Chapter 9 - Production of Dicarboxylic Acid Platform Chemicals Using Yeasts: Focus on Succinic Acid, p. 237–269. In Poltronieri, P, D’Urso, OF (eds.), Biotransformation of Agricultural Waste and By-Products. Elsevier.

25. Lazuka A, Auer L, Bozonnet S, Morgavi DP, O’Donohue M, Hernandez-Raquet G. 2015. Efficient anaerobic transformation of raw wheat straw by a robust cow rumen-derived microbial consortium. Bioresource technology 196:241–249.

26. Lewis NE, Nagarajan H, Palsson BO. 2012. Constraining the metabolic genotype–phenotype relationship using a phylogeny of in silico methods. Nature Reviews Microbiology 10:291–305.

27. Fang X, Lloyd CJ, Palsson BO. 2020. Reconstructing organisms in silico: genome-scale models and their emerging applications. Nat Rev Microbiol 18:731–743.

28. El-Semman IE, Karlsson FH, Shoaie S, Nookaew I, Soliman TH, Nielsen J. 2014. Genome-scale metabolic reconstructions of *Bifidobacterium adolescentis* L2-32 and *Faecalibacterium prausnitzii* A2-165 and their interaction. BMC Systems Biology 8:1–11.

29. Heinken A, Khan MT, Paglia G, Rodionov DA, Harmsen HJM, Thiele I. 2014. Functional metabolic map of *Faecalibacterium prausnitzii*, a beneficial human gut microbe. Journal of Bacteriology 196:3289–3302.

30. Magnúsdóttir S, Heinken A, Kutt L, Ravcheev DA, Bauer E, Noronha A, Greenhalgh K, Jäger C, Baginska J, Wilmes P, Fleming RMT, Thiele I. 2017. Generation of genome-scale metabolic reconstructions for 773 members of the human gut microbiota. Nature Biotechnology 35:81–89.

31. Bannink A, Lingen HJ van, Ellis JL, France J, Dijkstra J. 2016. The contribution of mathematical modeling to understanding dynamic aspects of rumen metabolism. Frontiers in Microbiology 7:1–16.

32. Hungate RE, Smith W, Clarke RTJ. 1966. Suitability of Butyl Rubber Stoppers for Closing Anaerobic Roll Culture Tubes. J Bacteriol 91:908–909.

33. Maglione G, Russell JB, Wilson DB. 1997. Kinetics of Cellulose Digestion by *Fibrobacter succinogenes* S85. Applied and Environmental Microbiology 63:665 LP – 669.

34. Miller GL. 1959. Use of Dinitrosalicylic Acid Reagent for Determination of Reducing Sugar. Analytical Chemistry 31:426–428.

35. Aite M, Chevallier M, Frioux C, Trottier C, Got J, Cortés MP, Mendoza SN, Carrier G, Dameron O, Guillaudeux N, Latorre M, Loira N, Markov GV, Maass A, Siegel A. 2018. Traceability, reproducibility and wiki-exploration for “à-la-carte” reconstructions of genome-scale metabolic models. PLoS Computational Biology 14:1–25.

36. Karp PD, Paley S, Romero P. 2002. The Pathway Tools software. Bioinformatics 18:S225–S232.

37. Karp PD, Billington R, Caspi R, Fulcher CA, Latendresse M, Kothari A, Keseler IM, Krummenacker M, Midford PE, Ong Q, Ong WK, Paley SM, Subhraveti P. 2019. The BioCyc collection of microbial genomes and metabolic pathways. Briefings in Bioinformatics 20:1085–1093.

38. Emms DM, Kelly S. 2015. OrthoFinder: solving fundamental biases in whole genome comparisons dramatically improves orthogroup inference accuracy. Genome Biol 16:157.

39. Emms DM, Kelly S. 2019. OrthoFinder: phylogenetic orthology inference for comparative genomics. Genome Biol 20:238.

40. Prigent S, Frioux C, Dittami SM, Thiele S, Larhlimi A, Collet G, Gutknecht F, Got J, Eveillard D, Bourdon J, Plewniak F, Tonon T, Siegel A. 2017. Meneco, a topology-based Gap-filling tool applicable to degraded genome-wide metabolic networks. Plos Computational Biology 13:e1005276.

41. Ebrahim A, Lerman JA, Palsson BO, Hyduke DR. 2013. COBRApy: COnstraints-Based Reconstruction and Analysis for Python. BMC Syst Biol 7:74.

42. Orth JD, Conrad TM, Na J, Lerman JA, Nam H, Feist AM, Palsson B. 2011. A comprehensive genome-scale reconstruction of *Escherichia coli* metabolism-2011. Molecular Systems Biology 7:535.

43. Heinken A, Sahoo S, Fleming RMT, Thiele I. 2013. Systems-level characterization of a host-microbe metabolic symbiosis in the mammalian gut. Gut Microbes 4:28–40.

44. Teusink B, Wiersma A, Molenaar D, Francke C, Vos WMD, Siezen RJ, Smid EJ. 2006. Analysis of growth of *Lactobacillus plantarum* WCFS1 on a complex medium using a genome-scale metabolic model. Journal of Biological Chemistry 281:40041–40048.

45. Caspi R, Billington R, Keseler IM, Kothari A, Krummenacker M, Midford PE, Ong WK, Paley S, Subhraveti P, Karp PD. 2020. The MetaCyc database of metabolic pathways and enzymes - a 2019 update. Nucleic Acids Research 48:D445–D453.

46. Moretti S, Martin O, Van Du Tran T, Bridge A, Morgat A, Pagni M. 2016. MetaNetX/MNXref – reconciliation of metabolites and biochemical reactions to bring together genome-scale metabolic networks. Nucleic Acids Res 44:D523–D526.

47. Belcour A, Girard J, Aite M, Delage L, Trottier C, Marteau C, Leroux C, Dittami SM, Sauleau P, Corre E, Nicolas J, Boyen C, Leblanc C, Collén J, Siegel A, Markov GV. 2020. Inferring biochemical reactions and metabolite structures to understand metabolic pathway drift. iScience 23:100849.

48. Altschul SF, Gish W, Miller W, Myers EW, Lipman DJ. 1990. Basic local alignment search tool. Journal of Molecular Biology 215:403–410.

49. Provost A, Bastin G. 2004. Dynamic metabolic modelling under the balanced growth condition. Journal of Process Control 14:717–728.

50. Provost A, Bastin G, Agathos SN, Schneider Y-J. 2006. Metabolic design of macroscopic bioreaction models: application to Chinese hamster ovary cells. Bioprocess Biosyst Eng 29:349–366.

51. Terzer M, Stelling J. 2008. Large-scale computation of elementary flux modes with bit pattern trees. Bioinformatics (Oxford, England) 24:2229–35.

52. Klamt S, Saez-Rodriguez J, Gilles ED. 2007. Structural and functional analysis of cellular networks with CellNetAnalyzer. BMC Systems Biology 1:2.

53. Song H-S, Ramkrishna D. 2009. Reduction of a set of elementary modes using yield analysis. Biotechnology and bioengineering 102:554–68.

54. Van Klinken JB, Willems Van Dijk K. 2016. FluxModeCalculator: an efficient tool for large-scale flux mode computation. Bioinformatics 32:1265–1266.

55. Singh D, Lercher MJ. 2020. Network reduction methods for genome-scale metabolic models. Cellular and Molecular Life Sciences 77:481–488.

56. Lugar DJ, Mack SG, Sriram G. 2021. NetRed, an algorithm to reduce genome-scale metabolic networks and facilitate the analysis of flux predictions. Metabolic Engineering 65:207–222.

57. Heirendt L, Arreckx S, Pfau T, Mendoza SN, Richelle A, Heinken A, Haraldsdóttir HS, Wachowiak J, Keating SM, Vlasov V, Magnusdóttir S, Ng CY, Preciat G, Žagare A, Chan SHJ, Aurich MK, Clancy CM, Modamio J, Sauls JT, Noronha A, Bordbar A, Cousins B, El Assal DC, Valcarcel LV, Apaolaza I, Ghaderi S, Ahookhosh M, Ben Guebila M, Kostromins A, Sompairac N, Le HM, Ma D, Sun Y, Wang L, Yurkovich JT, Oliveira MAP, Vuong PT, El Assal LP, Kuperstein I, Zinovyev A, Hinton HS, Bryant WA, Aragón Artacho FJ, Planes FJ, Stalidzans E, Maass A, Vempala S, Hucka M, Saunders MA, Maranas CD, Lewis NE, Sauter T, Palsson BØ, Thiele I, Fleming RMT. 2019. Creation and analysis of biochemical constraint-based models using the COBRA Toolbox v.3.0. Nat Protoc 14:639–702.

58. Muñoz-Tamayo R, Laroche B, Leclerc M, Walter E. 2009. IDEAS: A parameter identification toolbox with symbolic analysis of uncertainty and its application to biological modelling, p. 1271–1276. In IFAC Proceedings Volumes.

59. Muñoz-Tamayo R, Giger-Reverdin S, Sauvant D. 2016. Mechanistic modelling of in vitro fermentation and methane production by rumen microbiota. Animal Feed Science and Technology 220:1–21.

60. Vanrolleghem P. 1995. Practical identifiability of a biokinetic model of activated sludge respiration. Water Research 29:2561–2570.

61. Vavilin VA, Fernandez B, Palatsi J, Flotats X. 2008. Hydrolysis kinetics in anaerobic degradation of particulate organic material: An overview. Waste Management 28:939–951.

62. Muñoz-Tamayo R, Nielsen BL, Gagaoua M, Gondret F, Krause ET, Morgavi DP, Olsson IAS, Pastell M, Taghipoor M, Tedeschi L, Veissier I, Nawroth C. 2022. Seven steps to enhance Open Science practices in animal science. PNAS Nexus 1:pgac106.

63. Kanehisa M. 2000. KEGG: Kyoto Encyclopedia of Genes and Genomes. Nucleic Acids Research 28:27–30.

64. King ZA, Lu J, Dräger A, Miller P, Federowicz S, Lerman JA, Ebrahim A, Palsson BO, Lewis NE. 2016. BiGG Models: A platform for integrating, standardizing and sharing genome-scale models. Nucleic Acids Research 44:D515–D522.

65. Bansal P, Morgat A, Axelsen KB, Muthukrishnan V, Coudert E, Aimo L, Hyka-Nouspikel N, Gasteiger E, Kerhornou A, Neto TB, Pozzato M, Blatter M-C, Ignatchenko A, Redaschi N, Bridge A. 2022. Rhea, the reaction knowledgebase in 2022. Nucleic Acids Research 50:D693–D700.

66. Guiavarch E, Pons A, Creuly C, Dussap C-G. 2008. Application of a data reconciliation method to the stoichiometric analysis of *Fibrobacter succinogenes* growth. Appl Biochem Biotechnol 151:201–210.

67. Song H-S, Ramkrishna D. 2009. Reduction of a set of elementary modes using yield analysis. Biotechnology and bioengineering 102:554–68.

68. Robles-Rodriguez CE, Bideaux C, Guillouet SE, Gorret N, Cescut J, Uribelarrea JL, Molina-Jouve C, Roux G, Aceves-Lara CA. 2017. Dynamic metabolic modeling of lipid accumulation and citric acid production by *Yarrowia lipolytica*. Computers & Chemical Engineering 100:139–152.

69. Kass M, Ramin M, Hanigan MD, Huhtanen P. 2022. Comparison of Molly and Karoline models to predict methane production in growing and dairy cattle. Journal of Dairy Science 105:3049–3063.

70. Ellis JL, Dijkstra J, Kebreab E, Bannink A, Odongo NE, McBride BW, France J. 2008. Aspects of rumen microbiology central to mechanistic modelling of methane production in cattle. Journal of Agricultural Science 146:213–233.

71. Islam MM, Fernando SC, Saha R. 2019. Metabolic modeling elucidates the transactions in the rumen microbiome and the shifts upon virome interactions. Frontiers in Microbiology 10:1–17.

72. Lee NR, Lee CH, Lee DY, Park JB. 2020. Genome-scale metabolic network reconstruction and in silico analysis of hexanoic acid producing *Megasphaera elsdenii*. Microorganisms 8.

73. Agren R, Liu L, Shoaie S, Vongsangnak W, Nookaew I, Nielsen J. 2013. The RAVEN Toolbox and Its Use for Generating a Genome-scale Metabolic Model for *Penicillium chrysogenum*. PLoS Comput Biol 9.

74. Capela J, Lagoa D, Rodrigues R, Cunha E, Cruz F, Barbosa A, Bastos J, Lima D, Ferreira EC, Rocha M, Dias O. 2022. *merlin*, an improved framework for the reconstruction of high-quality genome-scale metabolic models. Nucleic Acids Research 50:6052–6066.

75. Arkin AP, Cottingham RW, Henry CS, Harris NL, Stevens RL, Maslov S, Dehal P, Ware D, Perez F, Canon S, Sneddon MW, Henderson ML, Riehl WJ, Murphy-Olson D, Chan SY, Kamimura RT, Kumari S, Drake MM, Brettin TS, Glass EM, Chivian D, Gunter D, Weston DJ, Allen BH, Baumohl J, Best AA, Bowen B, Brenner SE, Bun CC, Chandonia JM, Chia JM, Colasanti R, Conrad N, Davis JJ, Davison BH, Dejongh M, Devoid S, Dietrich E, Dubchak I, Edirisinghe JN, Fang G, Faria JP, Frybarger PM, Gerlach W, Gerstein M, Greiner A, Gurtowski J, Haun HL, He F, Jain R, Joachimiak MP, Keegan KP, Kondo S, Kumar V, Land ML, Meyer F, Mills M, Novichkov PS, Oh T, Olsen GJ, Olson R, Parrello B, Pasternak S, Pearson E, Poon SS, Price GA, Ramakrishnan S, Ranjan P, Ronald PC, Schatz MC, Seaver SMD, Shukla M, Sutormin RA, Syed MH, Thomason J, Tintle NL, Wang D, Xia F, Yoo H, Yoo S, Yu D. 2018. KBase: The United States department of energy systems biology knowledgebase. Nature Biotechnology 36:566–569.

76. Overbeek R, Disz T, Stevens R. 2004. The SEED: a peer-to-peer environment for genome annotation. Commun ACM 47:46–51.

77. Machado D, Andrejev S, Tramontano M, Patil KR. 2018. Fast automated reconstruction of genome-scale metabolic models for microbial species and communities. Nucleic Acids Research 46:7542–7553.

78. Zimmermann J, Kaleta C, Waschina S. 2021. gapseq: informed prediction of bacterial metabolic pathways and reconstruction of accurate metabolic models. Genome Biol 22:81.

79. Kanehisa M, Furumichi M, Tanabe M, Sato Y, Morishima K. 2017. KEGG: new perspectives on genomes, pathways, diseases and drugs. Nucleic Acids Res 45:D353–D361.

80. Henry CS, DeJongh M, Best A a, Frybarger PM, Linsay B, Stevens RL. 2010. High-throughput generation, optimization and analysis of genome-scale metabolic models. Nature biotechnology 28:977–82.

81. Mendoza SN, Olivier BG, Molenaar D, Teusink B. 2019. A systematic assessment of current genome-scale metabolic reconstruction tools. Genome Biol 20:158.

82. Schöpping M, Gaspar P, Neves AR, Franzén CJ, Zeidan AA. 2021. Identifying the essential nutritional requirements of the probiotic bacteria *Bifidobacterium animalis* and *Bifidobacterium longum* through genome-scale modeling. npj Syst Biol Appl 7:47.

83. Richards MA, Lie TJ, Zhang J, Ragsdale SW, Leigh JA, Price ND. 2016. Exploring hydrogenotrophic methanogenesis: A genome scale metabolic reconstruction of *Methanococcus maripaludis*. Journal of Bacteriology 198:3379–3390.

84. Zhu Y, Zhao J, Maifiah MHM, Velkov T, Schreiber F, Li J. 2019. Metabolic Responses to Polymyxin Treatment in *Acinetobacter baumannii* ATCC 19606: Integrating Transcriptomics and Metabolomics with Genome-Scale Metabolic Modeling. mSystems 4:1–15.

85. Venturelli OS, Carr AV, Fisher G, Hsu RH, Lau R, Bowen BP, Hromada S, Northen T, Arkin AP. 2018. Deciphering microbial interactions in synthetic human gut microbiome communities. Molecular Systems Biology 14:1–19.

86. Shoaie S, Karlsson F, Mardinoglu A, Nookaew I, Bordel S, Nielsen J. 2013. Understanding the interactions between bacteria in the human gut through metabolic modeling. Scientific Reports 3:1–10.

87. Levy R, Borenstein E. 2013. Metabolic modeling of species interaction in the human microbiome elucidates community-level assembly rules. Proceeding of the National Academy of Sciences of the United States of America 110:12804–12809.

88. Pereira B, Miguel J, Vilaça P, Soares S, Rocha I, Carneiro S. 2018. Reconstruction of a genome-scale metabolic model for *Actinobacillus succinogenes* 130Z. BMC Syst Biol 12:61.

89. Schuster S, Hilgetag C. 1994. On elementary flux modes in biochemical reaction systems at steady state. Journal of Biological Systems 2:165–182.

90. Singh D, Lercher MJ. 2020. Network reduction methods for genome-scale metabolic models. Cell Mol Life Sci 77:481–488.

91. Robles-Rodriguez CE, Bideaux C, Gaucel S, Laroche B, Gorret N, Aceves-Lara CA. 2014. Reduction of metabolic models by polygons optimization method applied to Bioethanol production with co-substrates. IFAC Proceedings 47:3: 6198–6203.

92. Muñoz-Tamayo R, Giger-Reverdin S, Sauvant D. 2016. Mechanistic modelling of in vitro fermentation and methane production by rumen microbiota. Animal Feed Science and Technology 220:1–21.

93. Matulova M, Delort AM, Nouaille R, Gaudet G, Forano E. 2001. Concurrent maltodextrin and cellodextrin synthesis by *Fibrobacter succinogenes* S85 as identified by 2D NMR spectroscopy. European Journal of Biochemistry 268:3907–3915.

94. Baroukh C, Muñoz-Tamayo R, Steyer JP, Bernard O. 2014. DRUM: A New Framework for Metabolic Modeling under Non-Balanced Growth. Application to the Carbon Metabolism of Unicellular Microalgae. PLoS One 9:e104499.

95. Verwoerd WS. 2011. A new computational method to split large biochemical networks into coherent subnets. BMC Systems Biology 5.

96. Schuster S, Pfeiffer T, Moldenhauer F, Koch I, Dandekar T. 2002. Exploring the pathway structure of metabolism: Decomposition into subnetworks and application to *Mycoplasma pneumoniae*. Bioinformatics 18:351–361.

97. Popova M, Fakih I, Forano E, Siegel A, Muñoz-Tamayo R, Morgavi DP. 2022. Rumen microbial genomics: from cells to genes (and back to cells). CABI Reviews https://doi.org/10.1079/cabireviews202217025.

