## Supplementary Material A for "Dynamic genome-based metabolic modeling of the predominant cellulolytic rumen bacterium *Fibrobacter succinogenes* S85"

**Table S1.** Compounds present in the biomass reaction of the large-scale metabolic network of *F. succinogenes* S85. The Table was built based on the biomass reaction modelled for *E. coli*.

| List of input compounds of biomass reaction | | | |
|---|---|---|---|
| **MetaCyc id** | **Name** | **Class** | **Coefficient** |
| ATP | ATP | nucleic acid | 40.1701 |
| CTP | CTP | nucleic acid | 0.12988 |
| DATP | dATP | nucleic acid | 0.023503 |
| DCTP | dCTP | nucleic acid | 0.023503 |
| DGTP | dGTP | nucleic acid | 0.023503 |
| GTP | GTP | nucleic acid | 0.2091 |
| TTP | dTTP | nucleic acid | 0.023503 |
| UTP | UTP | nucleic acid | 0.14027 |
| ARG | L-arginine | amino acid | 0.28828 |
| ASN | L-asparagine | amino acid | 0.23468 |
| CYS | L-cysteine | amino acid | 0.08898 |
| GLN | L-glutamine | amino acid | 0.25601 |
| GLT | L-glutamate | amino acid | 0.25601 |
| GLY | glycine | amino acid | 0.5958 |
| HIS | L-histidine | amino acid | 0.092623 |
| ILE | L-isoleucine | amino acid | 0.28255 |
| L-ALPHA-ALANINE | L-alanine | amino acid | 0.50006 |
| L-ASPARTATE | L-aspartate | amino acid | 0.23468 |
| LEU | L-leucine | amino acid | 0.43866 |
| LYS | L-lysine | amino acid | 0.33355 |
| MET | L-methionine | amino acid | 0.14934 |
| PHE | L-phenylalanine | amino acid | 0.18056 |
| PRO | L-proline | amino acid | 0.21543 |
| S-ADENOSYLMETHIONINE | S-adenosyl-L-methionine | amino acid | 0.004668 |
| SER | L-serine | amino acid | 0.2097 |
| THR | L-threonine | amino acid | 0.24665 |
| TRP | L-tryptophan | amino acid | 0.055157 |
| TYR | L-tyrosine | amino acid | 0.13425 |
| VAL | L-valine | amino acid | 0.4116 |
| SPERMIDINE | spermidine | amine | 0.004668 |
| CPD-9649 | all-trans-undecaprenyl diphosphate | lipid | 0.092476 |
| PROTOHEME | protoheme | organic heterocycle compound | 0.004668 |
| Glycogens | glycogen | carbohy-drates | 0.154187 |
| 5-METHYL-THF | 5-methyltetrahydropteroyl mono-L-glutamate | cofactor | 0.004668 |
| 10-FORMYL-THF | 10-formyl-tetrahydrofolate mono-L-glutamate | cofactor | 0.004668 |

| | | | |
|---|---|---|---|
| CO-A | coenzyme A | cofactor | 0.004668 |
| PYRIDOXAL_PHOSPHATE | pyridoxal 5’-phosphate | cofactor | 0.004668 |
| THF | 5,6,7,8-tetrahydrofolate | cofactor | 0.004668 |
| CPD-12125 | menaquinol-7 | acceptor | 0.004668 |
| FAD | FAD | acceptor | 0.004668 |
| NAD | $NAD^+$ | acceptor | 0.004668 |
| NADP | $NADP^+$ | acceptor | 0.004668 |
| REDUCED-MENAQUINONE | menaquinol-8 | acceptor | 0.004668 |
| CA+2 | $Ca^{2+}$ | ion | 0.004668 |
| CO+2 | $Co^{2+}$ | ion | 0.004668 |
| FE+2 | $Fe^{2+}$ | ion | 0.004668 |
| FE+3 | $Fe^{3+}$ | ion | 0.004668 |
| MG+2 | $Mg^{2+}$ | ion | 0.004668 |
| MN+2 | $Mn^{2+}$ | ion | 0.004668 |
| Na+ | $Na^+$ | ion | 0.004668 |
| ZN+2 | $Zn^{2+}$ | ion | 0.004668 |
| ADENOSYLCOBALAMIN | adenosylcobalamin | vitamin | 0.004668 |
| BIOTIN | biotin | vitamin | 0.004668 |
| RIBOFLAVIN | riboflavin | vitamin | 0.004668 |
| KCL | potassium chloride | salt | 0.004668 |
| WATER | $H_2O$ | | 34.7965 |
| **List of output compounds of biomass reaction** | | | |
| MetaCyc id | Name | Class | Coefficient |
| Bio | external biomass | | 1.0 |
| ADP | ADP | nucleic acid | 40.0 |
| Pi | phosphate | ion | 39.9953 |
| PPI | diphosphate | ion | 0.60238 |
| PROTON | $H^+$ | ion | 40.0 |

**Table S2.** Seeds and targets used for the GEM metabolic network reconstruction.

**Seeds**

| Name | Metacyc cpmpound Name | Id Metacyc |
|---|---|---|
| Cellulose | CELLULOSE | CELLULOSE |
| Cellobiose | Cellobiose | CELLOBIOSE |
| Glucose | Glucopyranose | Glucopyranose |
| Biotine | biotin | BIOTIN |
| p-aminobenzoic acid | 4-aminobenzoate | P-AMINO-BENZOATE |
| Ammonia (NH3) | ammonia | AMMONIUM |
| Water | water | WATER |
| Acetic acid | acetate | ACET |
| isobutyric acid | isobutanoate | ISOBUTYRATE |
| n-butyric acid | butanoate | BUTYRIC_ACID |
| DL-a-methyl butyric acid | 2-methylbutanoate | CPD-7076 |
| isovaleric acid | isovalerate | ISOVALERATE |
| n-valeric acid | valerianic acid | 5-AMINOPENTANOATE |
| propionic acid | propanoate | PROPIONATE |
| hemin | hemin | CPD-11678 |
| cysteine | L-cysteine | CYS |
| K2HPO4 | dipotassium phosphate | CPD0-2433 |
| MgSO4 | magnesium sulfate | CPD0-2390 |
| CaCl2 | calcium chloride | CPD0-1589 |
| MnSO4·6H2O | manganese sulfate | CPD0-1608 |
| CoCl2·6H2O | cobalt dichloride | CPD0-1695 |
| FeSO4·7H2O | iron sulfate | CPD0-2386 |
| ADP | ADP | ADP |
| ATP | ATP | ATP |
| Bio | | |
| CARBON-DIOXIDE | CARBON-DIOXIDE | CARBON-DIOXIDE |
| CL- | chloride | CL- |
| CA+2 | calcium(II) | CA+2 |
| CO+2 | cobalt(II) | CO+2 |
| CO-A | coenzyme A | CO-A |
| COB-I-ALAMIN | COB-I-ALAMIN | COB-I-ALAMIN |
| cob(II)inamide | cob(II)inamide | CPD-20903 |
| flavin adenine dinucleotide | FAD | FAD |
| FE+2 | iron(II) | FE+2 |
| FE+3 | iron(III) | FE+3 |
| flavin mononucleotide | flavin mononucleotide | FMN |
| potassium(I) | potassium(I) | K+ |
| potassium chloride | potassium chloride | KCL |
| MG+2 | magnesium(II) | MG+2 |
| MN+2 | manganese(II) | MN+2 |
| Na+ | SodiumI | Na+ |
| NACL | Sodium chloride | NACL |
| NAD | NAD | NAD |
| NADH | NADH | NADH |
| NADP | NADP | NADP |

| | | |
|---|---|---|
| NADPH | NADPH | NADPH |
| (NH4)2SO4 | ammonium sulfate | NH42SO4 |
| **Targets** | | |
| **Name** | **Metacyc cpmpound Name** | **Id Metacyc** |
| Acetic acid | acetate | ACET |
| Acetyl CoA | acetyl-CoA | ACETYL-COA |
| Alanine | L-alanine | L-ALPHA-ALANINE |
| Arginine | L-arginine | ARG |
| Asparagine | L-asparagine | ASN |
| Aspartate | L-aspartate | L-ASPARTATE |
| biotin | biotin | BIOTIN |
| Citrate | citrate | CIT |
| Cysteine | L-cysteine | CYS |
| Folate | folate | CPD-12826 |
| Formate | formate | FORMATE |
| Fructose-6-P, | β-D-fructofuranose 6-phosphate | FRUCTOSE-6P |
| Fructose-1-6-P | β-D-fructose 1,6-bisphosphate | FRUCTOSE-16-DIPHOSPHATE |
| Glyceraldehyde-3-P | D-glyceraldehyde-3-P | GAP |
| 1,3-PP Glycerate | 3-phospho-D-glyceroyl-phosphate | DPG |
| 3-P Glycerate | 3-phospho-D-glycerate | G3P |
| 2-P Glycerate | 2-phospho-D-glycerate | 2-PG |
| Glycine | glycine | GLY |
| Glycogen (futyle cycle : synthetized and degraded Gaudet) | a glycogen | Glycogens |
| Glutamate | L-glutamate | GLT |
| Glutamine | L-glutamine | GLN |
| Histidine | L-histidine | HIS |
| Isocitrate | D-threo-isocitrate | THREO-DS-ISO-CITRATE |
| Isoleucine | L-isoleucine | ILE |
| Leucine | L-leucine | LEU |
| Lysine | L-lysine | LYS |
| Malate | (S)-malate | MAL |
| Meso 2,6 diaminopimelic acid | meso-diaminopimelate | MESO-DIAMINOPIMELATE |
| Methionine | L-methionine | MET |
| Oxaloacetate | oxaloacetate | OXALACETIC_ACID |
| 2-oxoglutarate | 2-oxoglutarate | 2-KETOGLUTARATE |
| Pantothenate | (R)-pantothenate | PANTOTHENATE |
| Phenylalanine | L-phenylalanine | PHE |
| Phosphoenolpyruvate (PEP) | phosphoenolpyruvate | PHOSPHO-ENOL-PYRUVATE |
| Proline | L-proline | PRO |
| Pyruvate | pyruvate | PYRUVATE |
| Ribose-5-P | D-ribose 5-phosphate | RIBOSE-5P |
| Ribulose-5-P | D-ribulose 5-phosphate | RIBULOSE-5P |

| | | |
|---|---|---|
| Serine<br>Succinyl CoA | L-serine<br>succinyl-CoA | SER<br>SUC-COA |
| Succinate | succinate | SUC |
| Threonine | L-threonine | THR |
| Tryptophan | L-tryptophan | TRP |
| Tyrosine | L-tyrosine | TYR |
| Valine | L-valine | VAL |
| Xylulose-5-P | D-xylulose 5-phosphate | XYLULOSE-5-PHOSPHATE |
| UDP-N-acetyl-α-D-muramoyl-L-alanyl-γ-D-glutamyl-L-lysyl-D-alanyl-D-alanine | C3 | C3 |
| UDP-N-acetyl-α-D-muramoyl-L-alanyl-γ-D-glutamyl-meso-2,6-diaminopimeloyl-D-alanyl-D-alanine | C1 | C1 |
| UDP-N-acetyl-α-D-muramoyl-L-alanine | CPD0-1456 | CPD0-1456 |
| ditrans,octacis-undecaprenyldiphospho-N-acetyl-(N-acetyl-β-D-glucosaminyl)muramoyl-L-alanyl-γ-D-glutamyl-L-lysyl-D-alUDP-OHMYR-ACETYLGLUCOSAMINE anyl-D-alanine | CPD-7695 | CPD-7695 |
| UDP-N-acetyl-α-D-muramoyl-L-alanyl-γ-D-glutamyl-meso-2,6-diaminopimelate | UDP-AAGM-DIAMINOHEPTANEDIOATE | UDP-AAGM-DIAMINOHEPTANEDIOATE |
| Adénine | adenine | ADENINE |
| Guanine | guanine | GUANINE |
| Cytosine | cytosine | CYTOSINE |
| Thymine | thymine | THYMINE |
| DCTP | 2'-deoxycytidine-5'-triphosphate<br>deoxycytidine-triphosphate<br>deoxy-CTP | DCTP |
| TTP | thiamine thiothiazolone diphosphate | TTP |
| GTP | GTP | GTP |
| NADP | NADP | NADP |
| 5-METHYL-THF | 5-methyltetrahydropteroyl mono-L-glutamate | 5-METHYL-THF |
| PYRIDOXAL_PHOSPHATE | PYRIDOXAL_PHOSPHATE | PYRIDOXAL_PHOSPHATE |
| CO-A | coenzyme A | CO-A |
| SPERMIDINE | SPERMIDINE | SPERMIDINE |
| THF | Oxolane | CPD-24834 |

| | | |
|---|---|---|
| CTP | cytidine-triphosphate | CTP |
| CPD-9649 | all-trans-undecaprenyl diphosphate | CPD-9649 |
| FAD | flavin adenine dinucleotide | FAD |
| PROTOHEME | PROTOHEME | PROTOHEME |
| ATP | ATP | ATP |
| S-ADENOSYLMETHIONINE | S-ADENOSYLMETHIONINE | S-ADENOSYLMETHIONINE |
| DGTP | DGTP | DGTP |
| ADENOSYLCOBALAMIN | ADENOSYLCOBALAMIN | ADENOSYLCOBALAMIN |
| 10-FORMYL-THF | 10-FORMYL-THF | 10-FORMYL-THF |
| CPD-12125 | menaquinol-7 | CPD-12125 |
| UTP | UTP | UTP |
| Glutamates | Glutamates | Glutamates |
| ACP | ACP | ACP |
| NAD | NAD | NAD |
| RIBOFLAVIN | RIBOFLAVIN | RIBOFLAVIN |
| REDUCED-MENAQUINONE | REDUCED-MENAQUINONE | REDUCED-MENAQUINONE |
| DATP | deoxy-ATP | DATP |
| PPI | diphosphate | PPI |
| Pi | phosphate | Pi |
| PROTON | PROTON | PROTON |
| ADP | ADP | ADP |

**Table S3:** Comparison of the two draft models obtained via Pathway Tools from the two annotated *F. succinogenes* genomes NC_017448 and NC_013410

| **Common reactions** | |
|---|---|
| **common reactions found with Pathway Tools in NC_017448 and NC_013410 genomes** | **817** |
| **Specific reactions** | |
| **6 specific reactions to NC_013410** | **4 specific reactions to NC_017448** |
| 1.2.7.8-RXN (gene: FISUC_RS06185)<br>3.8.1.11-RXN (gene: FISUC_RS04815)<br>ALDOSE-1-EPIMERASE-RXN (gene: FISUC_RS03645)<br>ALDOSE1EPIM-RXN (gene: FISUC_RS03645)<br>RXN-6263 (gene: FISUC_RS04815)<br>RXN-6264 (gene: FISUC_RS04815) | RXN0-5190 (genes: FSU_RS06885 or FSU_RS05220)<br>RXN-17391 (genes: FSU_RS06885 or FSU_RS05220)<br>RXN-17392 (genes: FSU_RS06885 or FSU_RS05220)<br>RXN-17393 (genes: FSU_RS06885 or FSU_RS05220) |
